# CurT dosage quantitatively affects thylakoid structure and PSII performance in *Synechocystis* sp. PCC 6803

**DOI:** 10.64898/2026.04.22.720114

**Authors:** Matthias Ostermeier, Anne-Christin Pohland, Marcel Dann

**Author notes:** These authors contributed equally (Matthias Ostermeier, Anne-Christin Pohland). The authors responsible for distribution of materials integral to the findings presented in this article in accordance with the policy described in the Instructions for Authors are: Matthias Ostermeier and Marcel Dann.

## Abstract

The photosynthetic thylakoid membrane systems of many oxygenic photosynthesizers form elaborate three-dimensional structures. Curvature Thylakoid (CurT/CURT1) proteins are central regulators of thylakoid architecture in cyanobacteria and plants, driving both grana stacking and the formation of thylakoid convergence zones (TCZs) by promoting membrane curvature while also contributing to cyanobacterial cell and green-algal chloroplast division. While the functional role of grana stacks in terrestrial photosynthesis is well established, the physiological significance of cyanobacterial TCZs remains unclear. Here, we investigated the role of TCZs in *Synechocystis* sp. PCC 6803 by quantitatively assessing how CurT protein abundance shapes thylakoid membrane organization, TCZ frequency, and thylakoid layering, and how these architectural features relate to photosynthetic efficiency in a set of *curT* expression mutants. By correlating defined structural parameters of the thylakoid system with cellular CurT levels, we defined for the first time the quantitative relationship between CurT abundance, thylakoid architecture, and photosystem II (PSII) performance. Our results reveal a non-linear, logarithmic relationship: minimal CurT suffices to restore TCZ formation and sustain WT-like growth, while overexpression further enhances PSII activity and culture yield. Together, these findings identify CurT as a dose-dependent key determinant of thylakoid structure and underscore the functional importance of thylakoid membrane architecture for efficient photosynthesis.

**Significance Statement:** This study assesses how the membrane-shaping protein CurT quantitatively regulates thylakoid architecture and photosynthetic performance in the model cyanobacterium *Synechocystis* sp. PCC 6803. By relating CurT abundance to the formation of thylakoid convergence zones, photosystem II efficiency, and growth rate, we provide evidence of CurT acting as a key regulator of cyanobacterial membrane organization in a dosage-dependent manner. Our findings provide insight into the structural basis and physiological relevance of thylakoid convergence zones and suggest CurT as a potential target for improving cyanobacterial light-energy conversion.

## Introduction

Cyanobacteria are one of the most ancient and taxonomically diverse bacterial clades defined by their capacity for oxygenic photosynthesis (Garcia-Pichel et al., 2020), relying on the tandem operation of two light-driven membrane-embedded multi-protein redox enzyme complexes, *i.e.*, Photosystem II (PSII) and Photosystem I (PSI), to transfer electrons from water to ferredoxin *via* several intermediate electron carriers (Komenda et al., 2024; Qi et al., 2023).

The overall architecture of protein complexes driving the photosynthetic electron transport chain is highly conserved across cyanobacteria and is shared, with only minor modifications, by chloroplasts of eukaryotic algae and plants (Nickelsen & Rengstl, 2013). Apart from early-branching *Gloeobacterales* (Koyama et al., 2008; Rexroth et al., 2011), all these complexes are physically embedded in a specialized internal membrane system known as thylakoids (Ostermeier et al., 2024). Despite their conserved function, thylakoid systems show striking architectural diversity among oxygenic photosynthesizers (Mareš et al., 2019; Ostermeier et al., 2024). In land plant chloroplasts, thylakoids are commonly differentiated into loose stroma lamellae and tightly appressed and contorted grana stacks (Austin & Staehelin, 2011) which are considered an evolutionary adaptation to dry, high-light terrestrial environments (Gu et al., 2022). Meanwhile, cyanobacterial thylakoids are frequently arranged as concentric lamellae, parallel stacks, or irregular networks that pervade the cytoplasm, optimizing light capture and electron transport in low-light aquatic environments (Mareš et al., 2019). The precise architecture of these membrane systems varies markedly among species, reflecting adaptations to distinct ecological niches and cell morphologies.

*Synechocystis* sp. PCC 6803 (hereafter *Synechocystis*), a widely studied model organism, provides a prime example of specialized thylakoid architecture. In *Synechocystis*, thylakoids are organized into discrete peripheral layers or fascicles that lie just beneath the cytoplasmic membrane, leaving a central cytoplasmic core largely devoid of photosynthetic membranes (Heinz et al., 2016; Liberton et al., 2006; Ostermeier et al., 2024; Ostermeier et al., 2022; van de Meene et al., 2006). Within this peripheral zone, thylakoid convergence zones (TCZs) appear at specific sites where thylakoid membranes curve towards the plasma membrane (PM) into a compact junction before branching again into individual stacks and forming thylapses - in analogy to neural synapses – as close contact sites that likely facilitate lipid and small molecule exchange (Rast et al., 2019). This process may be aided by Vesicle-inducing protein in plastids 1 (VIPP1) (*aka* inner membrane-associated protein of 30 kDa IM30)-like proteins whose lipid-trafficking function is conserved even in *Gloeobacter* (Ma et al., 2025). The geometry of *Synechocystis* thylapse regions is often triangular or trapezoidal in crossLJsection, creating localized microdomains distinct from the more planar thylakoid lamellae (Rast et al., 2019). Recent studies have shown that two key membraneLJshaping factors, CurT and AncM (anchor of convergence membranes), act together to generate and stabilize TCZs in *Synechocystis* (Ostermeier et al., 2022). Here, CurT is the cyanobacterial homolog of CURVATURE THYLAKOID1 (CURT1) proteins first characterized in *Arabidopsis thaliana* (hereafter *Arabidopsis*) (Armbruster et al., 2013), and its activity is seemingly indispensable for forming the narrow, highly curved junctions at which thylakoid lamellae converge toward the plasma membrane (Heinz et al., 2016). In *Arabidopsis*, lossLJ and gainLJofLJfunction mutants demonstrated that CURT1 proteins oligomerize at grana margins and induce the extreme membrane curvature required for grana stacking (Armbruster et al., 2013; Pribil et al., 2018). Crucially, recombinant *Arabidopsis* CURT1 proteins suffice to tubulate liposomes *in vitro*, proving that membrane bending by CURT1 is an intrinsic, proteinLJmediated process (Armbruster et al., 2013).

In cyanobacteria, evidence points towards CurT fulfilling an analogous role. *Synechocystis* CurT tubulates liposomes *in vitro* (Heinz et al., 2016), can be functionally substituted by *Arabidopsis* CURT1A (Armbruster et al., 2013), and, while localizing broadly across thylakoid membranes, accumulates at TCZs (Heinz et al., 2016). CurT-deficient *Synechocystis* mutants completely lack organized convergence zones, and their thylakoid layers instead form disordered rings or crumpled arrays. This structural disruption coincides with a ∼50% reduction in PSII assembly (Heinz et al., 2016). LiveLJcell imaging further showed that CurT forms tubular networks at the periphery of the thylakoid system, consistent with a role in shaping membrane curvature at nascent convergence sites (Heinz et al., 2016; Ostermeier et al., 2022). Recently, in *Synechococcus elongatus* PCC 7942, CurT was shown to be also responsible for thylakoid architecture and photosynthetic performance (Zhang et al., 2025), thus corroborating a phylogenetically conserved role of CurT/CURT1-like proteins.

The AncM protein has been identified through a suppressor screen in a *curT*LJdeficient *Synechocystis* mutant (Ostermeier et al., 2022). This single-pass transmembrane protein localizes to punctate sites near the cell envelope, anchoring thylapse regions to the plasma membrane precisely where CurTLJinduced thylakoid curvature is most pronounced. In the absence of AncM, TCZs lose contact with the plasma membrane, resulting in enlarged lamellar stacks and diminished functional integrity (Ostermeier et al., 2022). Thus, CurT has been suggested to provide the membrane-bending force shaping the thylakoid lamellae, while AncM is hypothesized to secure curved membrane sectors in place, forming a structural scaffold that defines PSII assembly microdomains (Ostermeier et al., 2025; Ostermeier et al., 2022).

Structurally, CurT is assumed to oligomerize into highLJmolecularLJmass complexes that insert into the outer leaflet of the thylakoid bilayer, imposing local curvature that both concentrates and excludes specific lipid species (Heinz et al., 2016). The resulting curvature may promote spatial segregation of PSII assembly intermediates and create curvatureLJsensing niches that recruit additional biogenesis factors (Ostermeier et al., 2025; Rengstl et al., 2011). A sufficient degree of membrane curvature in turn could be essential for optimal phycobilisome-photosystem interaction and may also influence the contact of other soluble proteins with membrane-bound components (Lea-Smith et al., 2016). The precise molecular mechanisms and the extend to which such microdomain formation depends on protein-protein interactions, induced changes in lipid environments, and synergistic effects thereof, however, remain to be elucidated.

Previous studies have established that CurT deficiency affects both cell division and TCZ formation in cyanobacteria (Dann et al., 2025; Heinz et al., 2016; Ostermeier et al., 2022). Significant progress has also been made in uncovering the molecular foundation of TCZ and thylapse formation (Huokko et al., 2021; Ostermeier et al., 2024). In addition, TCZs have been functionally linked to PSII assembly (Ostermeier et al., 2025). However, the apparent absence of TCZ-like structures in many cyanobacterial taxa raises questions about their physiological relevance in *Synechocystis*. Moreover, the quantitative effects of CurT depletion and overproduction on photosynthesis remain uncharacterized. To address this, we analyzed a set of recently established *curT* expression level mutants (Dann et al., 2025) with elevated (overexpression, OE), reduced (complementation strain, compl.), and abolished (knock-out, KO) cellular CurT levels, as well as the parental wildtype (WT). We examined how relative CurT abundance influences thylakoid structural features measured as average numbers of TCZs and thylakoid layers per fascicle per cell, as well as photosynthetic efficiency assessed by PSII dark and light-adapted quantum yield and net oxygen evolution rates under photoautotrophic growth conditions. We found evidence of CurT depletion and enrichment having opposite effects on growth, PSII activity, and thylakoid organization: OE cells outperform WT, whereas compl. cells display intermediate phenotypes between KO and WT. Notably, the compl. strain revealed a non-linear relationship between CurT levels, TCZ formation, and PSII performance, indicating that even minimal amounts of cellular CurT are sufficient to support TCZ formation and cell division.

## Results and Discussion

### Cellular CurT levels affect PSII performance parameters under photoautotrophic growth conditions

A previously established set of *Synechocystis curT* expression mutants producing no (Δ*curT*_SpecR knockout mutant, KO), less (Δ*curT*_SpecR *slr0168*::*curT*_CmR complementation strain, compl.), or elevated (WT + *slr0168*::*curT*_SpecR overexpression mutant, OE) CurT was cultivated under photoautotrophic conditions at 25 °C and 50 µmol m^-2^ s^-1^ of warm-white LED light in BG11 medium until the mid-exponential growth phase was reached (see Methods). Samples obtained from these cultures were analyzed by transmission-electron microscopy (TEM) to assess cellular ultrastructure, by in-gel fluorescence assays and immunoblotting to quantify combined allophycocyanin+phycocyanin (APC+PC), CurT and PsbA (D1) protein levels, and by methanolic pigment extraction, oxygen evolution and chlorophyll fluorescence measurements to correlate thylakoid morphology, growth, and photosynthetic performance to cellular CurT levels.

Under photoautotrophic growth conditions, CurT depletion led to slower growth in KO cells, with average duplication times increasing by 57% (*P*<0.001) (**Fig. 1a, b**). The compl. strain showed a modest, non-significant increase in average duplication time (+12%, *P*=0.100) as compared to WT (**Fig. 1a, b**). In contrast, OE cells, displayed slightly shorter average duplication time (−4%; *P*=0.458) (**Fig. 1a, b**), which resulted in a significantly increased average OD_720_ at the end of the growth experiment (*i.e*., after 152 h of growth; +15%, *P*=0.007) (**Fig. 1c**). These results were in line with previous observations under photomixotrophic growth conditions (Dann et al., 2025; Heinz et al., 2016). Exponential-phase culture OD_750_ was found to closely correlate in a linear fashion with cellular dry weight (R^2^=0.95) (**Fig. 1d**), and cell count per unit OD_730_ was found virtually identical among all mutant strains (OE +1%; KO −2%; compl. −1%) (**Fig. 1e**). This finding corroborates previous reports on close correspondence between cell number and OD_750_ in parental *Synechocystis* WT and *curT* knockout mutants (3.9×10^7^ and 3.7×10^7^ cells ml^-1^ OD_750_^-1^) (Heinz et al., 2016), and *Synechocystis* OD_730/750_ and biomass in general (Myers et al., 2013), thus rendering OD-normalization suitable for subsequent immunoblot analyses and physiological measurements. Immunodetection of proteins in whole-cell protein extracts confirmed the absence of CurT in KO cells, while compl. and OE cells displayed significantly decreased (−97.9%, *P*=0.004) and enhanced (+77.3%, *P*=0.009) average cellular CurT levels, respectively (**Fig. 1 f, g**). This was largely in accordance with CurT levels previously observed in mature culture cells grown under photomixotrophic conditions (Dann et al., 2025; Heinz et al., 2016), albeit OE CurT levels being slightly higher than those in previous reports. Intriguingly, simultaneously detected D1 protein showed significantly reduced average cellular abundance in OE cells (−52.6%, *P*<0.001), KO (−74.1%, *P*<0.001), and compl. cells (−27.1%, *P*=0.001) as compared to WT (**Fig. 1f, g**). In-gel fluorescence analysis of combined APC and PC fluorescence signals revealed distinct accumulation patterns. KO cells exhibited strongly reduced average cellular APC+PC levels (−38%; *P*=0.013), thus corroborating previous results (Heinz et al., 2016), while both OE and compl. strains showed slightly but non-significantly increased average APC+PC levels relative to WT (OE: +16%, *P*=0.337; compl.: +18%, *P*=0.384) (**Fig. 1f, g**). Previously, PC depletion has been reported to follow complete mutational loss of PSII in *Synechocystis* (Kılıç et al., 2022). The lack of correspondence between cellular D1 and phycobiliprotein accumulation hence suggests that CurT-related alterations in PSII abundance may cause atypical peripheral antenna accumulation patterns.

**Figure 1.**
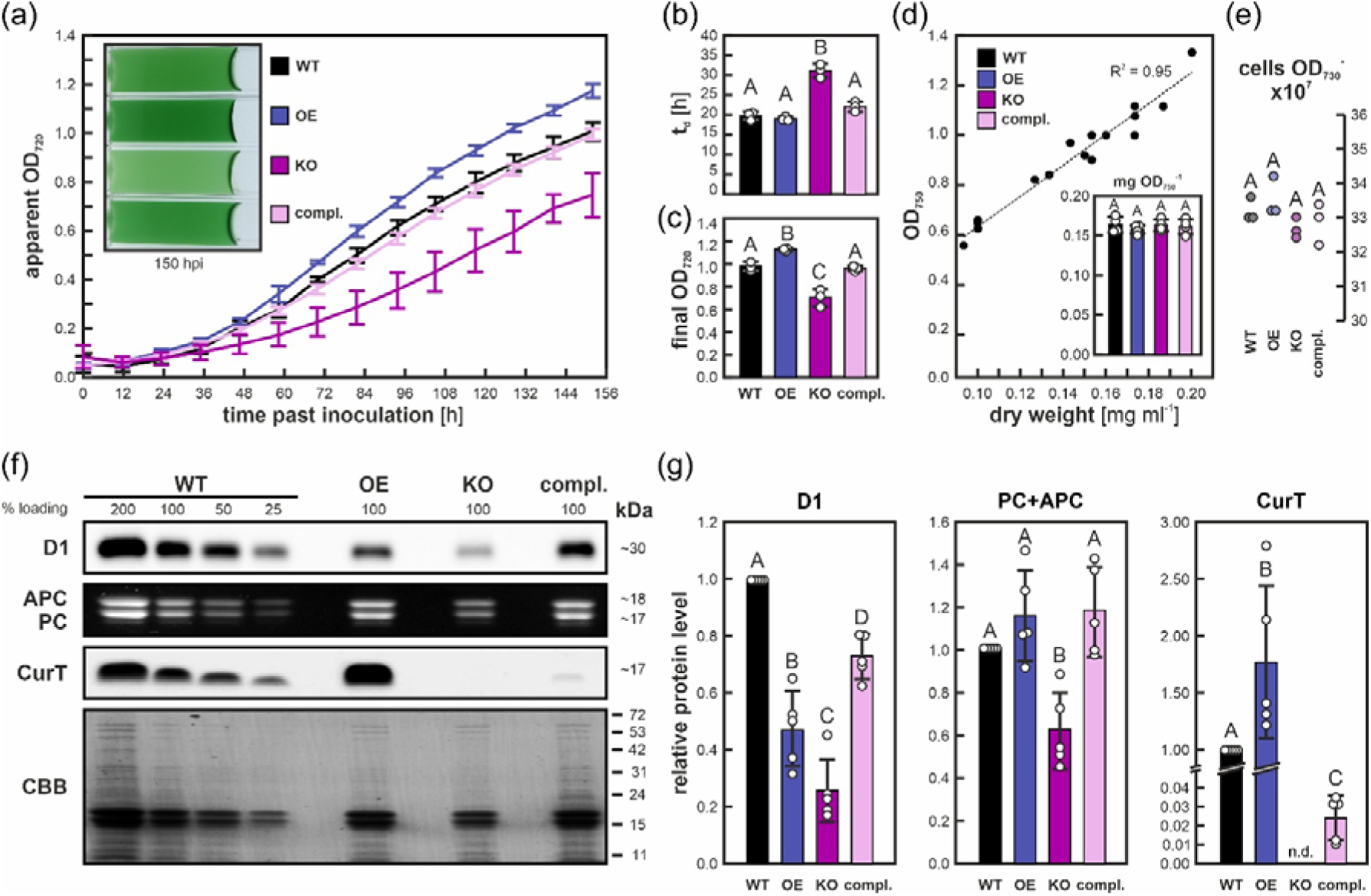
Photoautotrophic growth, CurT, and PSII marker protein level quantification in *Synechocystis curT* expression level mutants. (a) Growth curves of *curT* expression level mutants at 25 °C and 50 µmol photons m^−2^ s^−1^, monitored as apparent OD_720_ (Multicultivator inbuilt photometer; see Methods). Curves represent averages; error bars represent standard deviations of *n* = 3 biological replicates used for subsequent physiological characterization. Inset shows representative culture phenotypes at 150 hours past inoculation. WT, wildtype; OE, *curT* overexpression; KO, *curT* knock-out; compl., *curT* complementation strain. (b) Exponential growth phase duplication times t_d._ Columns show averages; error bars indicate standard deviations of *n* = 3 biological replicates. (c) Final apparent OD_720_ at 150 hours past inoculation. Columns show averages; error bars indicate standard deviations of *n* = 3 biological replicates. Individual data points are indicated. (d) Linear correlation between exponential-phase culture OD_750_ and dry weight of cell material. Inset shows corresponding dry-weight-to-unit-OD_750_ ratios. Columns show averages; error bars indicate standard deviations of *n* = 4 biological replicates. (e) Number of cells per unit OD_730_ as estimated by Neumann chamber count. Individual data points for *n* = 3 biological replicates are shown. (f) D1 and CurT protein levels as assessed by immunoblot analysis of whole-cell protein extracts obtained from mid-exponential phase cultures 3 days past inoculation. Combined allophycocyanin and phycocyanin (APC+PC) levels were assessed by in-gel fluorescent assay prior to blotting of SDS-PA gels. PVDF membrane Coomassie Brilliant Blue (CBB) staining served as loading control. Molecular weight markers are indicated. (g) Enhanced chemiluminescence and fluorescent signals were quantified to determine protein levels relative to 100% WT samples (see Methods). Data shown represents *n* = 5 biological replicates. Columns show averages; error bars indicate standard deviations. Individual data points are indicated. Uppercase letters indicate statistically significant differences (*p*<0.05) according to *post-hoc* Bonferroni-Holm-corrected Tukey HSD after significant among-group differences (*p*<0.05) had been determined by two-sided one-way ANOVA.

To scrutinize the apparent depletion of D1 on *curT* expression-level mutants, protein extracts derived from thylakoid membrane preparations were subjected to immunoblot analysis using Alexa Fluor 488-coupled secondary antibody. Blots with loading adjusted to chlorophyll *a* (Chl *a*) indicated no major decreases in D1, but showed mild yet consistent depletion in PsaA levels across all *curT* mutant strains (Supplementary Fig. 1a, b). Concurrently, ratios of quantitative PsaA and D1 protein level estimates (PsaA:D1) were found depleted to 57%, 89%, and 56% of the parental WT levels on average, respectively (Supplementary Fig. 1c). This difference in PSI to PSII was almost absent in low-temperature chlorophyll fluorescence spectrometry through comparison of emission spectra at 725 nm (PSI) and 695 nm (PSII) upon excitation with 438 nm wavelength, respectively, suggesting a mild relative depletion of PSI (F725:F695) in OE (−8%) and KO strains (−6%), but not compl. (−0%) (Supplementary Fig. 1d). As both immunoblot- and 77 K-derived PSI:PSII estimates are to be considered ambiguous at this point, precise effects of altered CurT levels on cellular PSI and PSII will have to be determined by quantitative proteomics (Jackson et al., 2023). Equally, possible effects of CurT abundance on PSI/PSII supercomplex formation, phycobilisome architecture, and state transitions affecting F725:F695-based PSI:PSII ratio estimates remain to be investigated (Acuña et al., 2018; Choubeh et al., 2018; Espinoza-Corral et al., 2024; Toyoshima et al., 2021; van Stokkum et al., 2023).

We next assessed the physiological effects of altered CurT abundance on PSII function and cellular pigment composition. Methanolic pigment extracts revealed that CurT depletion significantly increased abundance of cellular carotenoids (Car) relative to chlorophyll *a* (Chl*_a_*) in both KO (+43%, *P*<0.001) and compl. strains (+14%, *P*=0.006), but not OE (+4%, *P*=0.071) (Fig. 2 **a,c**), while absolute Chl*_a_* content per unit OD_750_ was unaltered in OE (−5%; *P*=0.187), decreased in KO (−24%; *P*<0.001) but increased in compl. (+11%; *P*=0.033) as compared to WT (Fig. 2b), corroborating previous observations from an independent *curT* deletion mutant (Heinz et al., 2016), while absolute carotenoid contents per unit OD_750_ were near-identical among all strains (Fig. 1b). Together this indicates changes in cellular CurT abundance and/or downstream effects thereof may quantitatively affect chlorophyll accumulation in *Synechocystis*. Combined with indication of changes in both absolute and relative photosystem content (Fig. 1g, Supplementary Fig. 1) and close correlation between OD_730/750_ with culture dry mass and cell count (Fig. 1d, e) across all strains, OD_750_-rather than Chl-*a*-normalization appears favorable for quantitative immunoblot analyses.

**Figure 2.**
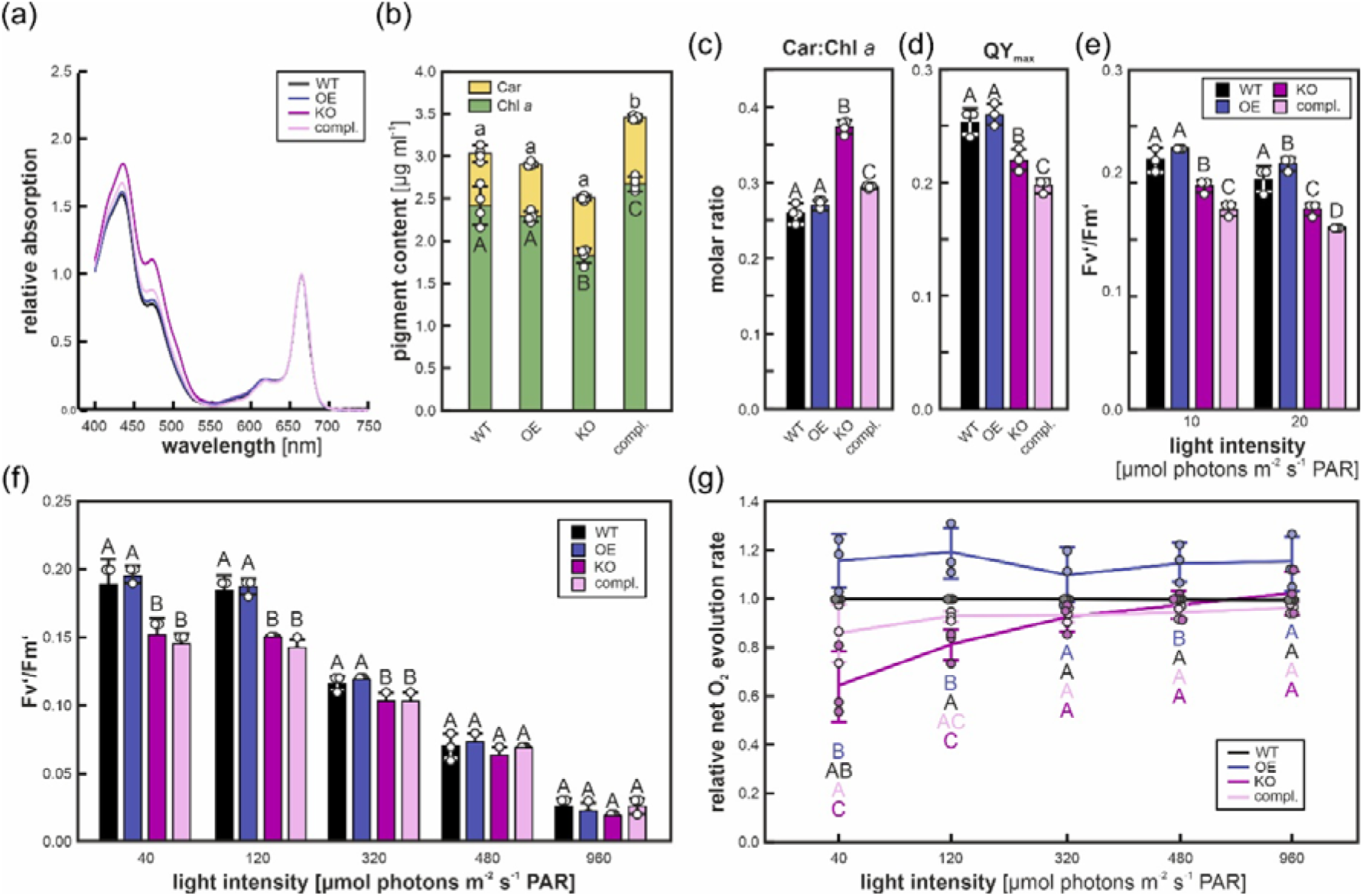
Cellular CurT protein levels affect pigmentation and PSII performance parameters in *Synechocystis*. (a) UV/Vis absorbance spectra of *Synechocystis* whole cell methanolic pigment extracts. Traces represent average absorbance spectra of *n* = 3 biological replicates normalized to the corresponding Q-band absorbance maxima. WT, wildtype; OE, *curT* overexpression; KO, *curT* knock-out; compl., *curT* complementation strain. (b) Chl *a* and carotenoid (Car) contents of OD_750_ = 0.75 cell equivalents. (c) Molar ratios of total carotenoids to chlorophyll *a* (Car:Chl*_a_*) in mid-exponential growth phase *Synechocystis curT* expression level mutants. (d) Apparent PSII maximum quantum yield (QY_max_) in mid-exponential growth phase dark-adapted *Synechocystis curT* expression level mutants. (e) PSII light-adapted quantum yields (Fv’/Fm’) in light-adapted mid-exponential growth phase *Synechocystis curT* expression level mutants exposed to low-light conditions. (f) Light-response curve of PSII light-adapted quantum yields (Fv’/Fm’) in light-adapted mid-exponential growth phase *Synechocystis curT* expression level mutants. (g) Light-response curve of net oxygen (O_2_) evolution rates per unit OD_750_ relative to WT measured in light-adapted mid-exponential growth phase *Synechocystis curT* expression level mutants at 25 °C.

PSII performance assays indicated impaired photochemistry in CurT-depleted strains, whereas OE cells performed slightly better than WT. Specifically, the maximum quantum yield of PSII of the dark-adapted samples (QY_max_) was significantly lower in both KO (−14%, *P*=0.008) and compl. (−22%, *P*<0.001) strains, while OE cells showed a slight, non-significant increase (+3%, *P*=0.419) (Fig. 2c). Despite physiological estimates of the quantum yield of PSII in cyanobacteria being quantitatively compromised in absence of herbicides (Ogawa et al., 2017), these results indicate a detrimental effect of CurT depletion or the differential loss of CurT-mediated thylakoid structural features on PSII functionality, possibly through impairment of *de-novo* biogenesis and/or the PSII repair cycle. Beyond independent quantitative assessment of CurT effects on cellular photosystem abundance and stoichiometry (see previous section; Supplementary Fig. 1), the precise identity and relative contributions of factors contributing to CurT-dosage-associated effects on PSII performance parameters require detailed future investigation. Such investigation will allow to dissect repair-specific and biogenesis-specific effects through, *e.g*., PSII-related complex profiling (Dobakova et al., 2007) and subcellular localization of PSII undergoing repair in hypothesized FtsH2-defined “repair zones” within the thylakoid system (Sacharz et al., 2015). The intriguing decline in QY_max_ of compl. mutant cells beyond that of fully segregated *curT* KO mutants may reflect an upregulation of compensatory mechanisms fostering PSII biogenesis occurring outside TCZs (Selao et al., 2016) in fully CurT-depleted cells, indicate a possible titration effect of TCZs on PSII assembly factors such as PratA (Stengel et al., 2012) and YCF48/HCF136 (Rengstl et al., 2011) on the rest of the cell which may disappear upon complete loss of TCZs, or hint at compensatory second-site mutations in the KO strain, as previously reported (Ostermeier et al., 2022), thus requiring further investigation of all three mechanisms.

Light-response curves obtained with an AquaPen fluorimeter showed that under low-light conditions (20 µmol photons m^-2^ s^-1^) PSII quantum efficiency in light-adapted state (F_v_’/F_m_’) was slightly yet significantly increased in OE cells (+7%, *P*=0.050) as compared to WT, while both KO (−13%, *P*=0.005) and compl. cells (−21%, *P*<0.001) showed significantly decreased F_v_’/F_m_’, respectively (Fig. 2d). At moderate light intensities (50-100 µmol photons m^-2^ s^-1^), KO cells initially displayed slightly higher F_v_’/F_m_’ values than compl. cells but dropped below them above 300 µmol photons m^-2^ s^-1^ (Fig. 2e). Meanwhile, both WT and OE PSII light-adapted quantum yield converged towards the lower levels seen in KO and compl. strains under high light (500-1000 µmol photons m^-2^ s^-1^) (Fig. 2e). This was paralleled by a stepwise convergence of relative net O_2_ evolution rates under increasing light intensities as measured with a Clark-type oxygen electrode. Here, at 40 µmol photons m^-2^ s^-1^, KO and compl. cell rates started out at 64% (*P*=0.020) and 81% (*P*=0.162) of WT and reached 103% (*P*=0.595) and 97% (*P*=1.184) at 960 µmol photons m^-2^ s^-1^, respectively. Meanwhile, OE cells displayed consistently increased net O_2_ evolution rates, starting at 116% (*P*=0.244) at 40 µmol photons m^-2^ s^-1^ and ending at 115% (*P*=0.162) of WT at 960 µmol photons m^-2^ s^-1^ (Fig. 2f). Importantly, the here-applied oxygen evolution measurement routine in absence of artificial PSII electron acceptors likely underestimates true PSII activity (Baikie et al., 2023; McCauley & Melis, 1987), but provides an informative measure for effective physiological OLJ output under variable light conditions (Dann et al., 2021). Due to technical limitations of the oxygen electrode and its inbuild illumination system (minimum light intensity 40 µmol photons m^-2^ s^-1^) used, no direct correlation could be drawn between low-light-adapted quantum yield estimates at 10 and 20 µmol photons m^-2^ s^-1^ and net oxygen evolution. Nevertheless, our observations indicate that higher CurT levels enhance PSII efficiency metrics (*i.e.*, QYmax and Fv’/Fm’) and enzymatic activity estimates (*i.e.*, net O_2_ evolution), particularly under low and moderate light intensities similar to the cultivation regime. Meanwhile, the elevated relative carotenoid content in CurT-depleted cells might reflects an increased demand for photoprotective pigments to mitigate oxidative stress during the PSII repair cycle (Izuhara et al., 2020; Kusama et al., 2015), but may also be attributed to a shift in cellular plasma-to-thylakoid membrane ratio as indicated by subsequent TEM analyses (Fig. 3) and previous reports (Heinz et al., 2016), thus increasing relative carotenoid contents due to high carotenoid abundance in the plasma membrane (Zhang et al., 2015). Still, as cellular levels of total carotenoids were not significantly altered in KO mutants (Fig. 2b), a higher relative carotenoid content within the less abundant thylakoid membranes of the KO mutant appears plausible.

**Figure 3.**
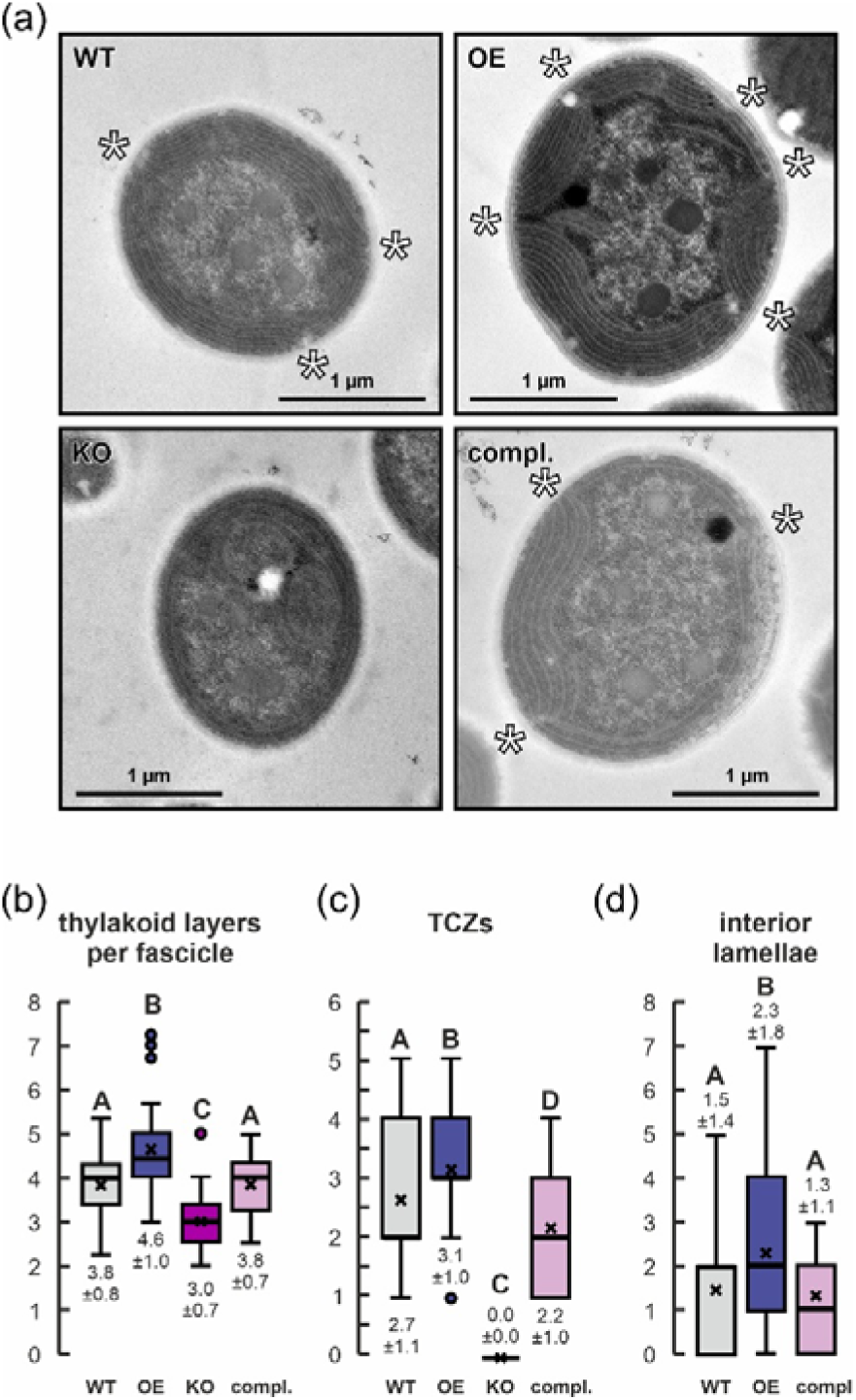
Dosage effects of cellular CurT levels on thylakoid system structure. (a) Representative transmission electron micrographs of *Synechocystis curT* expression level mutant cell thin sections. WT, wildtype; OE, *curT* overexpression; KO, *curT* knock-out; compl., *curT* complementation strain. White asterisks indicate thylakoid layers grouped to peripheral fascicle. Scale bars (1 µm) are indicated. (b) Average number of thylakoid layers per peripheral fascicle (*i.e.*, thylakoid-system segments interrupted by TCZs, see white asterisks in (a)) in mid-exponential growth phase *Synechocystis curT* expression level mutants. Note that in case of the KO mutant in absence of TCZs, this measure indicates the number of thylakoid layers per cell. (c) Number of thylakoid convergence zones in mid-exponential growth phase *Synechocystis curT* expression level mutants. (d) Number of thylakoid lamellae protruding into the cell center of mid-exponential growth phase *Synechocystis curT* expression level mutants.

In panels **b-e**, columns show averages; error bars indicate standard deviations of *n* = 3-4 biological replicates. Individual data points are indicated. Uppercase letters indicate statistically significant differences (*p*<0.05) according to *post-hoc* Bonferroni-Holm-corrected Tukey HSD after significant among-group differences (*p*<0.05) had been determined by two-sided one-way ANOVA.

Collectively, these results indicate that cellular CurT levels affect PSII peripheral antenna protein accumulation (Fig. 1f, g) and photochemical performance under photoautotrophic conditions (Fig. 2d**-g**), underpinning a quantitative role of CurT or CurT-mediated thylakoid structures on PSII assembly and/or repair in *Synechocystis*. CurT depletion or reduction (KO, compl.) appears to result in compromised PSII core abundance (Fig. 1f, g; but see also Supplementary Fig. 1), elevates relative cellular carotenoid content (Fig. 2a, c), slows growth (Fig. 1a, b), and lowers both PSII light-adapted quantum yield and net OLJ evolution (Fig. 2d**-g**). In contrast, CurT overexpression modestly enhances net OLJ evolution and late-stage culture density but also may disrupt cellular D1 accumulation, indicating that tight CurT homeostasis and CurT-related thylakoid structural features are critical for balanced PSII biogenesis and effective photoprotection.

### TCZ formation in Synechocystis quantitatively depends on CurT

Grana margins in plant chloroplasts and TCZs in cyanobacteria are highly curved thylakoid subdomains that concentrate the machinery required for photosystem assembly and repair (Ostermeier et al., 2025; Pribil et al., 2018). In *Arabidopsis*, the absence of CURT1 proteins results in chloroplasts containing flat, lobe-like thylakoids with considerably fewer grana margins and a decrease in photosynthetic performance; meanwhile, CURT1 overexpression increases grana stack height and therefore the number of grana margins, and significantly reduces grana stack diameter as compared with WT plants (Armbruster et al., 2013; Pribil et al., 2018). In *Synechocystis*, loss of CurT leads to a total loss of TCZs (Heinz et al., 2016), but the effects of both incomplete CurT depletion and enhanced CurT protein levels have not yet been studied in a quantitative manner. To this end, *curT* expression level mutants were subjected to TEM analysis, with cellular cross-sections being analysed as two-dimensional proxies for total three-dimensional cellular abundance of thylakoid system structural features.

TEM-micrographs of photoautotrophically grown *Synechocystis curT* expression level mutant cell thin-sections (Fig. 3a, Supplementary Fig. 3) revealed significant architectural differences among strains. OE cell thylakoid fascicles contained 21% thylakoid sheets on average (*P*<0.001), whereas the number of KO thylakoid membrane layers showed a significant decrease as compared to WT (−21%; *P*<0.001) (Fig. 3b). Meanwhile, the compl. strain did not show any obvious difference to WT (+0.1%; *P*=0.920) (Fig. 3b). The same cells showed a significant increase in the average number of TCZs in OE (+16%; *P*=0.031), while KO cells completely lacked (−100%; *P*<0.001) and compl. cells displayed a moderate yet statistically significant decrease in average numbers of TCZs per cell (−19%; *P*=0.024), respectively (Fig. 3c). These results indicate a quantitative effect of cellular CurT levels on both TCZ formation and thylakoid sheet number, complementing earlier qualitative observations on CurT being essential for TCZs formation (Heinz et al., 2016). Notably, this relationship appears to be non-linear as even trace amounts of cellular CurT in the compl. strain (∼2% of WT level; Fig. 1e) enable TCZ formation at approximately 79% the rate of WT cells (see Fig. 3c). This suggests that while CurT is essential for TCZ formation, most cellular CurT in WT is likely involved in other cellular processes or exists in an inactive protein reservoir. Additionally, OE cells displayed a significant increase in the average number of thylakoid lamellae protruding into the cell interior as compared to WT (+57%; *P*=0.031), while compl. cells showed a slight, non-significant reduction (−9%; *P*=0.707) (Fig. 3d). Such protruding lamellae may enhance the efficiency of RNA-binding proteins (RBPs) at thylakoid membranes oriented toward the cell interior, enabling the capture of psbA mRNA (Mahbub et al. (2020)). These membranes could also facilitate psbA mRNA transport and, ultimately, the insertion of the D1 polypeptide and PSII biogenesis at the TCZs (Ostermeier et al., 2025), thus potentially contributing to the enhanced late-stage culture density of OE mutant cells. In KO cells, no such lamellae could be observed, while vesicle-shaped thylakoid segments occupied the inside of most cells, consistent with previous findings (Heinz et al., 2016).

In panels **b-d**, boxplots represent data of *n* = 37/38/37/35 individual cells for WT/OE/KO/compl., respectively. Estimates are derived from one thin-section per observed cell. Boxplots centre lines indicate medians, crosses indicate averages, boxes indicate the 25th–75th percentiles, whiskers indicate the 1.5-fold interquartile range, and circles indicate outliers. Values below plots indicate averages ±LJstandard deviation. Uppercase letters indicate statistically significant differences (*p*<0.05) according to *post-hoc* Bonferroni-Holm-corrected Tukey HSD after significant among-group differences (*p*<0.05) had been determined by two-sided one-way ANOVA.

To quantify the relationship between CurT, thylakoid architecture, and photosynthetic performance, we correlated relative CurT protein levels (Fig. 1e) with structural and physiological parameters using logarithmic regression models (Fig. 4). Both the abundance of cellular TCZs and net O_2_ evolution rate under low-light conditions approximating the cultivation regime (*i.e.*, 40 µmol photon m^-2^ s^-1^) showed close logarithmic correlation with cellular CurT levels (R^2^ = 0.99 for TCZs; R^2^ = 0.89 for net O_2_; Fig. 4a, b), characterized by a steep increase in both TCZ formation and O_2_ evolution upon injection of trace amounts of CurT into the system. Likewise, a strong logarithmic correlation between CurT levels and exponential growth phase cell duplication time was observed (R^2^=0.99) (Fig. 4c), corroborating previous and current observations of low levels of cellular CurT facilitating efficient cell division under both photomixotrophic (Dann et al., 2025) and photoautotrophic growth conditions (Fig. 1a, b). Importantly, our findings indicate that CurT functions as a low-dosage architectural determinant of thylakoid organization and a facilitator of cell division but argue against its large-scale structural involvement in shaping thylakoid architecture. Instead, other yet-to-be identified protein factors may be involved. Alternatively, CurT-dependent recruitment of non-bilayer-forming galactolipids (Böde et al., 2025; Murphy, 1982) or induced changes in lipid density (de Jesus et al., 2013) may play a crucial role in promoting and stabilizing localized thylakoid membrane curvature, which may in turn mediate the recruitment of other specific protein components (Vanni et al., 2014). Meanwhile, increases in cellular CurT levels enhance both, number of thylakoid layers per peripheral fascicles and cellular growth, albeit at diminishing returns as indicated by the flat plateau phases common to all three logarithmic regressions (Fig. 3b**, 4**). WT CurT abundance therefore likely represents a physiological optimum under natural growth conditions, while artificial elevation of CurT levels may improve biomass yield in controlled culture systems as indicated by increased late-culture optical density and robust OD_750_-to-dry-mass correlation (Fig. 1c, d), suggesting potential for biotechnological application.

**Figure 4.**
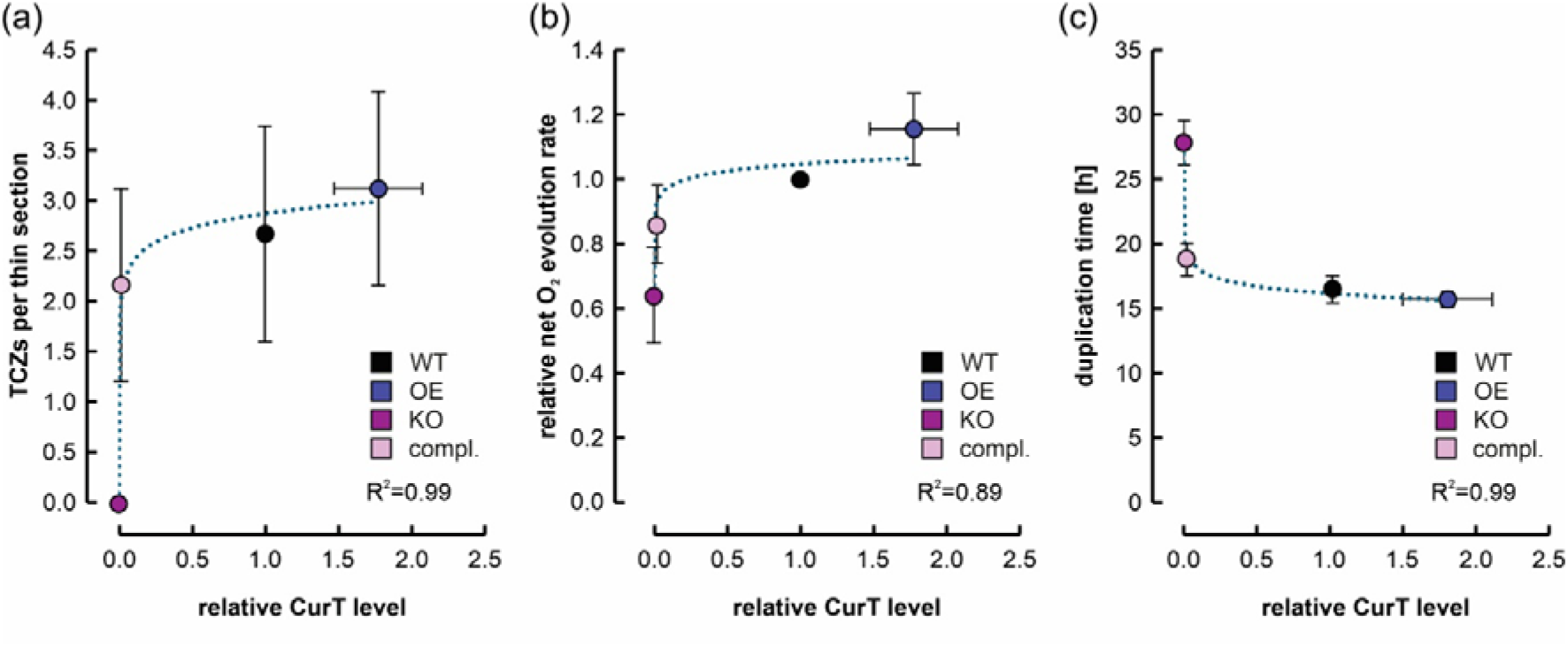
Thylakoid structural features, PSII activity, and cell division rates are a logarithmic function of *Synechocystis* cellular CurT protein levels. (a) Correlation of cellular CurT levels as determined by immunoblot analysis (Fig. 1e) with the number of thylakoid convergence zones (TCZs) per thin section of mid-exponential growth phase *Synechocystis curT* expression level mutant cells (Fig. 4c). Logarithmic trend function: y = 0.2071ln(x) + 2.8816. (b) Correlation of cellular CurT levels as determined by immunoblot analysis (Fig. 1e) with net oxygen (O_2_) evolution rates per unit OD_750_ relative to WT measured in light-adapted mid-exponential growth phase *Synechocystis curT* expression level mutants at 40 µmol photons m^-2^ s^-1^ and 25 °C, closely approximating cultivation conditions (Fig. 2f). Logarithmic trend function: y = 0.0309ln(x) + 1.0471. (c) Correlation of cellular CurT levels as determined by immunoblot analysis (Fig. 1e) with duplication times of mid-exponential growth phase *Synechocystis curT* expression level mutant cells (Fig. 1b). Logarithmic trend function: y = −0.833ln(x) + 19.411.

In panels **a-c**, coloured circles indicate average values, and error bars indicate standard deviations (vertical: TCZs/relative net O_2_ evolution rate/duplication time; *n* = 3) or standard errors (horizontal: relative cellular CurT levels; *n* = 5), respectively. Coefficients of determination R^2^ for logarithmic trends (dotted lines) are provided within each graph.

The observations outlined above grant quantitative insight into the relationship between cellular CurT abundance, thylakoid architecture, and physiology, pointing towards significant differences in photosynthetic efficiency arising from otherwise well-tolerated, CurT-level related changes in cellular ultrastructure. Remarkably, OE strains exhibit superior photosynthetic performance based on PSII performance parameters (Fig. 2d**-g**) and late-stage culture density (Fig. 1c) despite immunoblot data suggesting diminished cellular D1 levels (Fig. 1e). In contrast, compl. strains with similar indication of D1 depletion and a similar cellular phycobiliprotein content as OE showed impaired PSII performance (Fig. 2d**-g**) and increased relative carotenoid accumulation (Fig. 2c). A plausible explanation for this discrepancy lies in a reduced cellular frequency of TCZs in CurT-deficient compl. cells, possibly lengthening the diffusive pathways for PSII complexes undergoing repair. Recent findings show predominant localization of pre-PsbA2/3 to TCZs of *Synechocystis* (Ostermeier et al., 2025), suggesting these zones may serve not only as sites of *de-novo* PSII biogenesis, but also for the PSII repair cycle. Thus, decreases in PSII light-adapted quantum yield and net oxygen evolution in compl. cells may result from compromised trafficking of damaged PSII to the cellular repair sites. Future studies will have to directly trace PSII biogenesis and repair in order to elucidate CurT-dosage effects on both processes.

Finally, enhanced net O_2_ evolution and robust growth of OE cells despite lower cellular PSII levels may further stem from improved light capture due to a more homogenous distribution of thylakoid membranes throughout the cytoplasm (Fig. 5a). Specifically, the increased number of thylakoid lamellae protruding into the cell interior (Fig. 3d, Fig. 5b) may – in addition to a possible increase in the efficiency of RBP recruitment (Mahbub et al., 2020) – expand the effective light-absorption cross-section while at the same time reducing self-shading under low and moderate light intensities as indicated by increased PSII light-adapted quantum yields (Fig. 2d) and net O_2_ evolution rates (Fig. 2f) in OE cells. Importantly, the oxygen electrode setup used in this study has a lower limit of light intensity at 40 µmol photons m^-2^ s^-1^, necessitating further experiments to determine exact low-light performance parameters and light compensation points for OE and *curT*-depleted cells.

**Figure 5.**
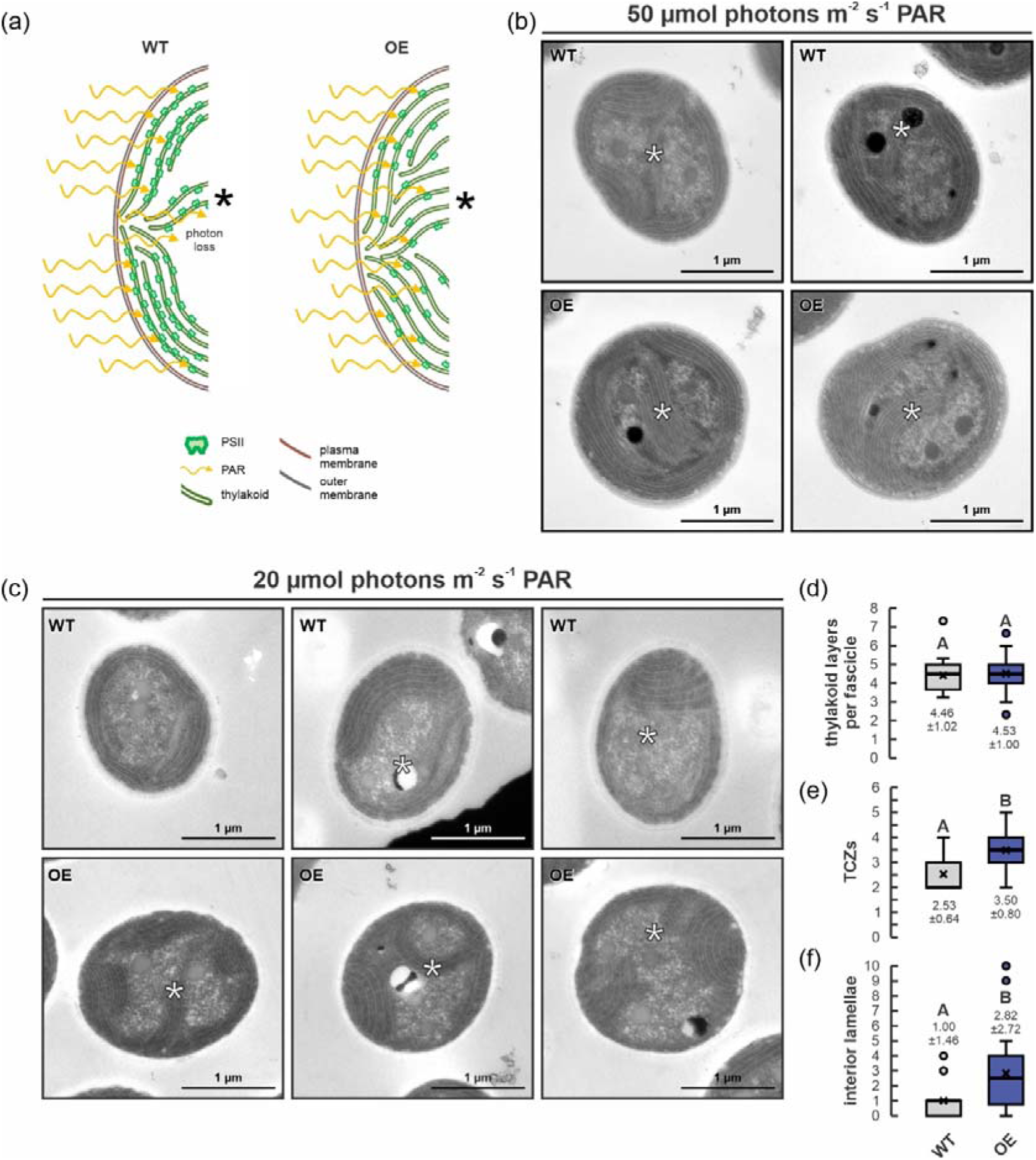
Structural basis for enhanced PSII performance parameters in *Synechocystis curT* overexpression mutants despite low cellular PSII. (a) Schematic representation of thylakoid structure and PSII distribution throughout the cell in wildtype (WT) and *curT* overexpression (OE) mutant cells. Comparably high density of PSII presumably leads to self-shading under low-light conditions, while PSII-depleted thylakoid convergence zones reduce local photon absorption efficiency. Additional thylakoid layers and lamellae protruding into the cell center may counteract these shortcomings in OE cells, resulting in overall more even PSII distribution throughout the cell, improving photon absorption and utilization. PAR, photosynthetically active radiation. Asterisks indicate interior thylakoid lamellae. (b) Representative thin sections transmission electron micrographs of *Synechocystis* WT and *curT* OE mutant cells grown under 50 µmol photons m^-2^ s^-1^ PAR illustrate increased numbers of thylakoid lamellae protruding into the center of OE cells (see also Fig. 3d). Scale bars (1 µm) are indicated. Asterisks indicate interior thylakoid lamellae. (c) Representative thin sections transmission electron micrographs of *Synechocystis* WT and *curT* OE mutant cells grown under 20 µmol photons m^-2^ s^-1^ PAR illustrate increased numbers of thylakoid lamellae protruding into the center of OE cells (white asteriks). Scale bars 1 µm. (d) Average number of thylakoid layers per peripheral fascicle (*i.e.*, thylakoid-system segments interrupted by TCZs, see white asterisks in (**a, c**)) in mid-exponential growth phase *Synechocystis* WT and *curT* OE mutant cells. (e) Number of thylakoid convergence zones in mid-exponential growth phase WT and *curT* OE mutant cells. (f) Number of thylakoid lamellae protruding into the cell center of mid-exponential growth phase WT and *curT* OE mutant cells.

CurT OE mutant cells grown under low-light intensity (20 µmol photons m^-2^ s^-1^ PAR) displayed similar ultrastructural changes as under standard light intensity (50 µmol photons m^-2^ s^-1^ PAR), with low-light OE cells showing significant increase in average number of TCZs (+38%; *P*<0.001) and interior thylakoid lamellae (+180%; *P*=0.013) as compared to WT (Fig. 5c**-f**), indicating a stable qualitative effect on thylakoid structure due to *curT* overexpression under reduced photon flux density. Meanwhile, under elevated photon flux density (300 µmol photons m^-2^ s^-1^ PAR), no discernable differences were observed between WT and OE thylakoid structure (Supplementary Fig. 4). In both cases a high amount of glycogen granula between the thylakoid membranes – likely a result of enhanced photosynthetic activity under high light – was shown, with large accumulations commonly overlapping with tentative TCZs, thus distorting the TEM signal and obstructing quantitative analysis. Meanwhile, CurT protein levels were found unaltered in WT cells grown under both low (109±17%; *P*=0.594) and high-light (118±22%; *P*=0.895) as compared to moderate light controls (Supplementary Fig. 5). This may indicate per-cell CurT accumulation to be largely irresponsive to fluctuations in light intensity within the assayed range of 20-300 µmol photons m^-2^ s^-1^ PAR and is in line with previous *Synechocystis* proteomics studies that did not find CurT (Slr0483) to be differentially expressed under 300 µmol photons m^-2^ s^-1^ PAR high-light conditions (Hong et al., 2014). However, previous microarray hybridization studies tracing *Synechocystis* transcriptome changes upon transition from 50 to 500 µmol photons m^-2^ s^-1^ PAR found *curT* mRNA significantly depleted under high light (Georg et al., 2009), pointing towards a possible depletion of cellular CurT beyond the here-tested high-light regime and rendering further specifics of light-dependency of CurT activity to be elucidated. Equally, the precise physiological effects of altered thylakoid structure under low light require further examination.

Notably, reduced self-shading through mutational truncation of peripheral antenna has previously been shown to enhance growth in both plants (Kirst et al., 2017) and cyanobacteria (Sengupta et al., 2023), but to our knowledge, no such effect has yet been reported to result from changes in thylakoid architecture and a reduction of PSII core antenna abundance as indicated for CurT OE strains. Here, additional protruding thylakoid lamellae may facilitate light harvesting in CurT-over-accumulating cells, possibly compensating for reductions in cellular PSII abundance to some degree. Although correlative light and electron microscopy (CELM) has confirmed the photosynthetic activity of such internal thylakoid membranes (Lübben et al., 2024; Ostermeier et al., 2022), their precise functional contribution and possible side effects of their overabundance remain unsolved. Future studies should therefore investigate how altered thylakoid distribution and curvature affect overall photosynthetic efficiency in OE cells, as well as the physiological constraints that may limit CurT accumulation in WT under natural conditions.

In panels d-f, boxplots represent data of *n* = 15/22 individual cells for WT/OE, respectively. Estimates are derived from one thin-section per observed cell. Boxplots centre lines indicate medians, crosses indicate averages, boxes indicate the 25th–75th percentiles, whiskers indicate the 1.5-fold interquartile range, and circles indicate outliers. Values below plots indicate averages ±LJstandard deviation. Uppercase letters indicate statistically significant differences (p<0.05) according to two-sided *t*-test.

## Conclusion

In conclusion, our results establish CurT as a dosage dependent key regulator of thylakoid architecture, and photosynthetic performance in *Synechocystis*. Controlled modulation of cellular CurT levels produced predictable changes in thylakoid sheet number and TCZ abundance, both of which correlated strongly with PSII assembly, light-adapted quantum yield, and net oxygen evolution. Remarkably, trace amounts of CurT (∼2% of WT) were sufficient to restore the majority of TCZ formation (∼79% of WT), revealing a non-linear threshold behavior in which minimal CurT fulfils essential biogenic functions, while higher levels further promote cellular thylakoid membrane accumulation (as evidenced by increased numbers of thylakoid layers per peripheral fascicle and cell-interior thylakoid lamellae) and final culture density. CurT overexpression increased net OLJ evolution and final culture yield but seemingly disrupted D1 accumulation, emphasizing the importance of tight CurT homeostasis for balanced PSII biogenesis and photoprotection. Together, these findings suggest that CurT-induced thylakoid restructuring and provision of PSII-assembly microdomains functionally benefits photosynthesis under low and moderate light intensities. With recent evidence pointing towards partial conservation of CurT functionality in thylakoid and PSII biogenesis (Zhang et al., 2025) and previous reports of thylakoid lamellae extending toward the plasma membrane during early thylakoid biogenesis in thylapse-like structures (Huokko et al., 2021) in *Synechococcus elongatus* PCC 7942, broader applicability of our findings to other cyanobacteria lacking typical TCZs appears plausible.

Importantly, our results indicate that TCZs are not merely structural features but have clear physiological significance: CurT-dependent TCZ formation correlates with PSII assembly/repair efficiency, net OLJ-evolution rates, and growth. This supports a model in which TCZs act as microdomains that concentrate assembly intermediates - non-exclusive but preferential - and shorten diffusion pathways *via* thylapses, thereby enhancing photosynthetic performance. However, because excessive CurT or the structures is induces appear to perturb D1 accumulation, TCZ biogenesis appears to be a tunable but tightly regulated determinant of thylakoid function. Future studies should hence elucidate the molecular mechanisms of CurT-mediated recruitment of biogenesis factors and assess its functional dynamics under diverse environmental conditions.

## Materials and Methods

### Cyanobacterial strains, culture conditions, and growth curve collection

Experiments were conducted with glucose-tolerant *Synechocystis* sp. PCC 6803 strains as used in (Dann et al., 2025). The mutant set comprised *curT* knock-out mutants with the *curT* coding sequence (ORF slr0483) replaced by a spectinomycin resistance marker (KO), overexpression mutants with an additional *curT* gene copy inserted into the genomic neutral site slr0168 (OE), and complementation strains carrying both the KO and OE mutant alleles. All mutant alleles were shown to be fully segregated (Dann et al., 2025). For maintenance, cultures were routinely grown under continuous illumination with 40 μmol photons m^-2^ s^-1^ white fluorescent light (shaker cultures: 5000 K, FHF32EX-N-HX-S; plate cultures: 4000 K; MonotaRO Co. Ltd., Hyogo, Japan) at 25 °C. For growth on solid media, BG11 was supplemented with 0.75% (w/v) bacteriological agar, 1 mM TES/KOH (pH 8.4), and 4 g L^-1^ sodium thiosulfate. Pre-cultures were grown with suitable antibiotics added to the culture medium (KO, OE: 25 µg ml^-1^ spectinomycin; compl.: 25 µg ml^-1^ spectinomycin 8.5 µg ml^-1^ chloramphenicol), while assay cultures were grown without antibiotics to minimize physiological stress. Assay liquid cultures were inoculated at OD_730_ = 0.05 in BG11 photoautotrophic medium (Rippka et al., 1979) and cultivated in a Multi-Cultivator MC 1000-OD (Photon Systems Instruments, Drásov, Czech Republic) under 50 µmol photons m^-2^ s^-1^ of warm-white LED light and atmospheric aeration, collecting growth curves in 60-min intervals using the built-in MC 1000-OD photometer. In the late exponential and early plateau phase, corresponding OD_720_ values collected from undiluted cultures represent underestimates of the “true” OD_720_ as measured in conventional 10 mm cuvettes due to the lack of dilution and thickness of the culture layer assayed (27 mm diameter of culture vessels), which is why these values are labelled “apparent OD_720_”. Assay culture samples for biochemical, physiological, and microscopic analyses were collected within the exponential growth phase, and total cultivation time approximately 150-160 h. For low- and high-light experiments, cells were grown under 20 and 300 µmol photons m^-2^ s^-1^, respectively, under otherwise identical conditions. *Synechocystis* strains were grown at 25 °C in order to preserve direct comparability with previous studies on the same mutant material (Dann et al., 2025).

*Synechocystis* duplication time estimates were calculated after the exponential growth phase was determined on a logarithmic scale plot for all strains as t_d_=LN(2)/([LN(OD_t1_)/LN(OD_t0_)]/[_t1_-_t0_]). Exponential growth phases were observed to occur between 29 (t_0_) and 56 (t_1_) hours past inoculation for WT, OE, and compl., and between 40 (t_0_) and 64 (t_1_) hours past inoculation for KO mutant strains.

### Synechocystis dry weight determination

Synechocystis exponential growth phase culture dry weight was determined by harvesting cell material from 15 ml culture volume by centrifugation (10 min, 3,200 x*g*), washing cells with 1 ml of BG11 media, and subsequently drying cell pellets completely in a rotary evaporator. Dry weight was then related to original OD_750_ determined for the corresponding cultures.

### Determination of Synechocystis cell count per unit OD_730_

To determine the cell count per unit OD_730_ in *Synechocystis curT* expression-level mutants, cells of exponential growth phase cultures were harvested and adjusted to OD_730_=0.8 and subsequently counted in a Neubauer-improved counting chamber (Carl Roth, Karlsruhe, Germany) under a lightmicroscope ZEISS Axioskop (ZEISS, Oberkochen, Germany).

### Protein extraction, detection, and quantification

*Synechocystis* whole-cell protein extracts were prepared as described by Dann et al. (2021) from cells grown in liquid culture for approximately 3 days to mid-exponential growth phase.

Culture samples were normalized to OD_750_, harvested by centrifugation, dissolving the cells in lysis buffer (Tris 100 mM - pH 6.8 (HCl); glycerol 24 % (v/v); SDS 5 % (w/v); bromophenol blue 0.02 % (w/v); DTT 100 mM) at 95 °C for 5 min under vigorous agitation (1200 rpm). Equivalents of OD_750_=4.0 cells were lysed in 100 µl buffer and debris was excluded by centrifugation prior to loading. For standard SDS PAGE, 5 µl of sample corresponding to OD_750_=0.2 cell equivalents were loaded. *Synechocystis* thylakoid membranes were prepared as previously described (Gandini et al., 2017) and sample concentration was adjusted to Chl *a* content. 10 μl of whole-cell protein extract or 10 µg Chl *a* equivalent thylakoid protein samples were fractionated by SDS PAGE on 4% /12% polyacrylamide Tris-Tricine gels (Schägger & Von Jagow, 1987), Proteins were blotted onto PVDF membrane (Millipore Immobilon-P Transfer Membrane, pore size 0.45 μm) at 0.1 A, 25 V for 90 min (Trans-Blot^®^ Turbo^TM^ Transfer System, Bio-Rad Laboratories, Inc., Hercules, CA, USA), and the CurT protein was immuno-detected using primary antibody serum raised against the N-terminus of *Synechocystis* CurT (Armbruster et al., 2013) provided by Dario Leister (LMU Munich) at a 1:10000 dilution. PsbA/D1 and PsaA proteins were immuno-detected using polyclonal primary antibody AS05 084 (D1) and AS06 172 (PsaA) (Agrisera, Sweden). Combined allophycocyanin and phycocyanin levels (APC+PC) were quantified *via* in-gel fluorescence using a BIO RAD ChemiDoc MP Imaging System (Alexa 546 fluorescence channel) prior to gel blotting. Immunoblot ECL signals were detected using horse-radish-peroxidase conjugated Goat anti-Rabbit IgG AS09 602 (Agrisera, Sweden) as secondary antibody at 1:10000 dilution. Immunoblot fluorescent signals were detected using Alexa FluorTM 488 goat anti-rabbit IgG (H+L) secondary antibody (Invitrogen A-11008). Chemiluminescence and fluorescence signals were quantified using ImageJ (Schneider et al., 2012), and normalized to the intensity of the corresponding wildtype signal on the respective PVDF membrane. As not all blots used for whole-cell protein level quantification (Fig. 1g, h) included a full WT titration series, protein level approximation was achieved through normalization to 100% WT signals on the respective blots. ECL signal intensities detected for the blot shown in Fig. 1g indicate a robust linear correlation between loading input and ECL signal intensity, however, supporting linear approximation (R^2^ 0.95-0.99; Supplementary Fig 2a). Further effective protein quantity estimates of thylakoid PsaA, D1, and CurT based on Alexa Fluor 488 fluorescence signals (Supplementary Fig 1) and ECL (Supplementary Fig 5) were derived using individual linear or logarithmic calibration curves for each blot and protein, with R^2^ values ranging from 0.93 to 0.99 (Supplementary Fig 2b, c; Supplementary Fig 6).

### Low-temperature chlorophyll fluorescence spectrometry (77 K)

The 77K fluorescence emission spectra of *Synechocystis* cells were measured using a HORIBA Fluoromax Plus FL-1013 (HORIBA Jobin Yvon GmbH, Bensheim, Germany) after chlorophyll excitation at 438 nm, adapted to Klinkert et al. (2004).

### Methanolic pigment extraction and quantification

Pigments were extracted and quantified according to the method described by Zavřel et al. (2015). For pigment analysis, pigments from 1 mL of *Synechocystis* culture with an OD_750_ of 0.75 were pelleted at 9,680 x*g* for four minutes. The pelleted cells were then resuspended in 1 mL of cold methanol, vortexed, and covered with aluminium foil. Pigments were extracted for four hours in the refrigerator. The solution was centrifuged at 9680 x*g* for 10 minutes, and the supernatant was decanted into a 10 mm polystyrene cuvette (Sarstedt, Nümbrecht, Germany), and the absorption spectrum was measured between 350 nm and 800 nm (FastGene^®^ Photometer NanoSpec (FG-NP01), NIPPON Genetics EUROPE GmbH, Düren, Germany).

### Fluorescence analysis to estimate maximum and light-adapted quantum yields

To estimate the performance of PSII in *Synechocystis* cells, fluorescence light curves were obtained using an AquaPen (AquaPen AP110-C, Photon Systems Instruments, Drásov, Czech Republic). The measuring pulses were set to 10%, and the saturating light pulses were set to 50%. To estimate the maximum (QYmax) and light-adapted quantum yield (Fv’/Fm’) under different light intensities, the cells were dark-adapted for ten minutes before initiating the predefined light curve three (LC3; light intensities in μmol photons m^-2^ s^-1^: 10, 20, 50, 100, 300, 500, 1000).

### O_2_ evolution measurements

Oxygen evolution measurements of *Synechocystis* cultures were performed as previously described (Dann et al., 2021). To analyze O_2_ evolution at PSII, ∼ 2 mL of cell culture was filled into the chamber of a Clark-Type Electrode Oxytherm+P (Hansatech Instruments Ltd., Pentney, Norfolk, United Kingdom). Measurements were conducted at 25 °C with light intensity changing every five minutes. The seven light intensity steps in μmol photons m^-2^ s^-1^ were: 0, 40, 120, 320, 480, 960, 0. Net O_2_ evolution was determined by correcting for the respiration rate obtained after the light regimen, and the data were normalized to the cultures’

OD_750_. To compare results among three sets of measurements obtained from separate electrode preparations and calibrations, results of the OE, KO and compl. strains were normalised to the corresponding WT controls.

### Transmission electron microscopy

For *Synechocystis* cells, sample preparation was performed as previously described (Lübben et al., 2024). Synechocystis cells were pelleted by centrifugation at 2,000 x*g* for 15 minutes. The resulting pellets were gently resuspended in 20 μl of culture medium immediately before cryofixation. Samples were frozen under high pressure (2,100 bar) using an EM HPM100 (Leica Microsystems, Wetzlar, Germany) and then stored in liquid nitrogen. Cryofixation was followed by freeze-substitution in A.O.U.H. solution at –90 °C (acetone containing 0.2% [w/v] OsOLJ, 0.1% [w/v] uranyl acetate, and 9% [v/v] H_2_O) for 42 hours, as previously described (Peschke et al., 2013; Walther & Ziegler, 2002), using an EM AFS2 (Leica Microsystems). After substitution, samples were embedded in Epon 812 and polymerized for 16 hours at 63 °C. Ultrathin sections of 70 nm (ultra 35°, 3.0 mm, DiATOME) were cut on an Ultracut E ultramicrotome (Leica Microsystems) using diamond knives. Sections were collected on collodion-coated, 400-mesh copper grids (Science Services GmbH, Munich, Germany). Prior to imaging, ultrathin sections were post-stained with lead citrate according to Reynolds (1963). Samples were examined in a Zeiss EM 912 transmission electron microscope (Zeiss, Oberkochen, Germany) equipped with an integrated OMEGA energy filter operated in zero-loss mode at 80 kV. Images were captured at a nominal magnification of 12,500× using a 2k × 2k slow-scan CCD camera (Tröndle Restlichtverstärkersystem, Moorenweis, Germany).

### Statistical Analyses

Charts were created using Microsoft Office Excel 365. For boxplots, internal datapoints are indicated, horizontal lines represent the median, crosses represent average values, and boxes indicate the 25th and 75th percentiles. Whiskers extend 1.5-fold the interquartile range with outliers being represented as circles beyond the range of the whiskers. Statistically significant among-group differences were tested for by one-way ANOVA (two-sided), followed by post-hoc Tukey HSD (honest significant differences) tests with Bonferroni–Holm p-value correction for multiple comparison. Significant differences according to multiple simultaneous post hoc comparisons are routinely indicated by uppercase letters denoting the resultant groups of not significantly different (same letter) and significantly different (different letter) samples. Analyses were performed using the one-way ANOVA with post-hoc test tool as implemented by Navendu Vasavada (https://astatsa.com/).

## Acknowledgements

We thank Ms Michaela Finke for photo-documentation of the *Synechocystis* cultures. Further we thank Prof. Dr. Jörg Nickelsen for basic support, Ms Jennifer Grünert for support in EM microscopy, and Dr Marta Ludwiczak and Prof. Dr. Dario Leister for support with the 77K measurements. This work was supported by grant DA2816/1-1 awarded to M.D. by the Deutsche Forschungsgemeinschaft.

## Author contributions

M.O. and M.D. designed the research; M.O., A.-C.P. and M.D. performed research; M.O., A.-C.P. and M.D. analyzed data; M.O. and M.D. wrote the manuscript with contributions from A.-C.P.

## Supplementary Figures

**Supplementary Figure 1.**
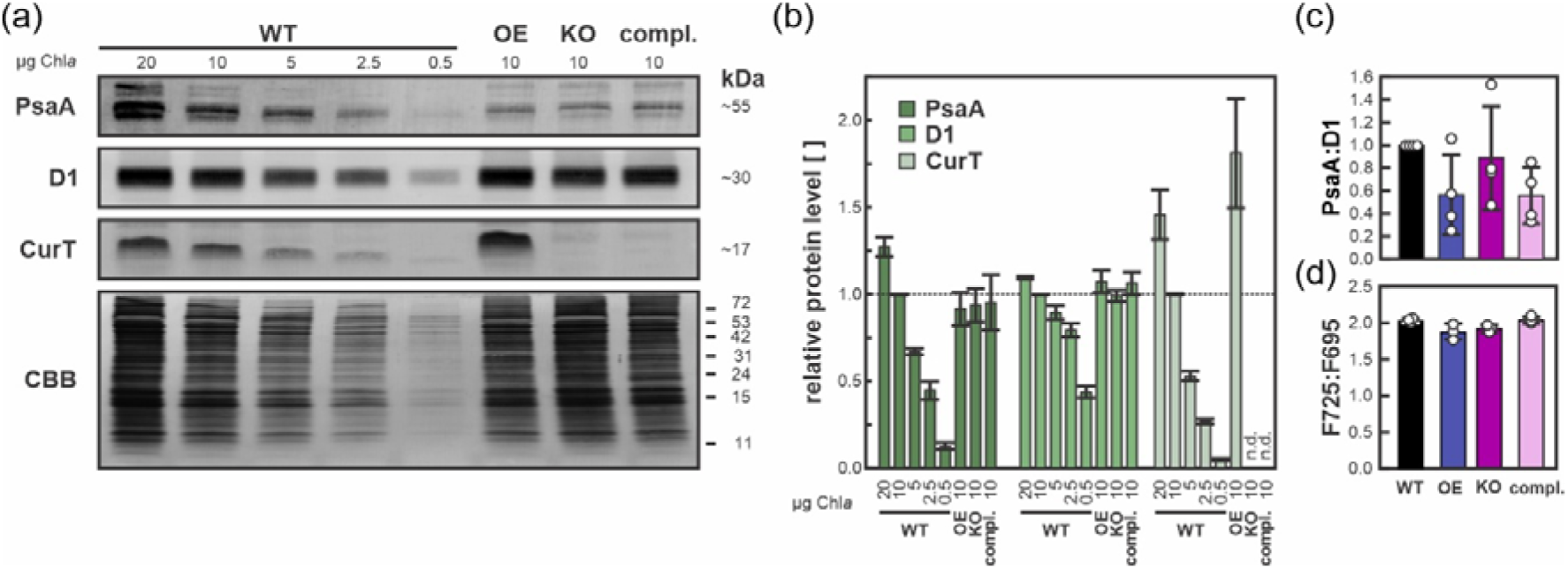
Protein-level indication of altered PSI:PSII ratio in *curT* expression mutants. (a) PsaA, D1, and CurT protein levels as assessed by immunoblot analysis of thylakoid-membrane preparation protein extracts obtained from mid-exponential phase cultures 3 days past inoculation. Loading was adjusted to Chl _a_ levels with 10 µg Chl *a* per lane as standard. PVDF membrane Coomassie Brilliant Blue (CBB) staining served as loading control. Molecular weight markers are indicated. (b) Secondary-antibody-conjugated Alexa Fluor 488 fluorescent signals relative to respective 10 µg Chl *a* WT samples. Data shown represents *n* = 4 biological replicates. Columns show averages; error bars indicate standard errors. Due to increased background signal as compared to ECL detection, CurT could not be detected in the complementation strain. (c) PsaA-to-D1 quantified protein signal ratios (see Methods). Data shown represents *n* = 4 biological replicates. Columns show averages; error bars indicate standard deviations. (d) PSI-to-PSII ratios derived from low-temperature chlorophyll fluorescence spectra recorded at 77 K. PSI- and PSII-specific emission was detected at 725 and 695 nm (F725 and F695), respectively. Data shown represents *n* = 3 biological replicates. Columns show averages; error bars indicate standard deviations. Individual data points are indicated.

**Supplementary Figure 2.**
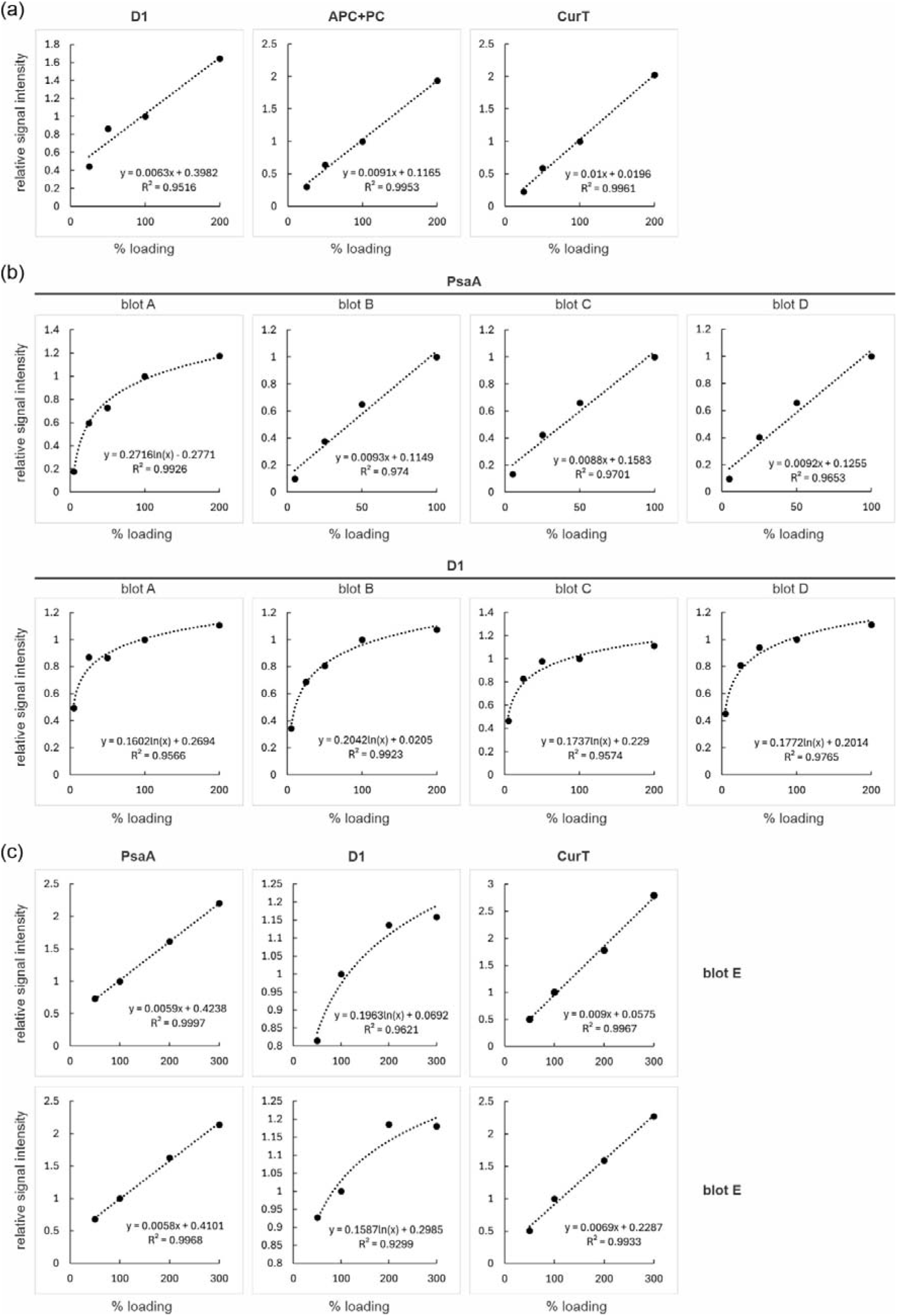
Calibration curves for *curT* expression mutant immunoblot signal quantification. (a) Exemplary calibration curves derived from immunoblot ECL signals shown in Fig. 1f indicate linear correlation of whole-cell protein sample input (x-axis; 100% corresponding to c) and ECL signal intensity relative to 100% input (y-axis). (b) Calibration curves derived from immunoblot Alexa Fluor 488 fluorescence signals shown in Supplementary Fig. 1a (plus three more biological replicates; blots A-D) indicate partially linear, partially logarithmic correlation of thylakoid-membrane preparation protein sample input (x-axis; 100% corresponding to 10 µg of Chl *a*) and Alexa Fluor 488 emission signal intensity relative to 100% input (y-axis). Best-fitting linear or logarithmic regression models were chosen covering the required range for *curT* OE, KO, and compl. signal quantification. Regression curve equations and corresponding coefficient of determination R^2^ are indicated.

**Supplementary Figure 3.**
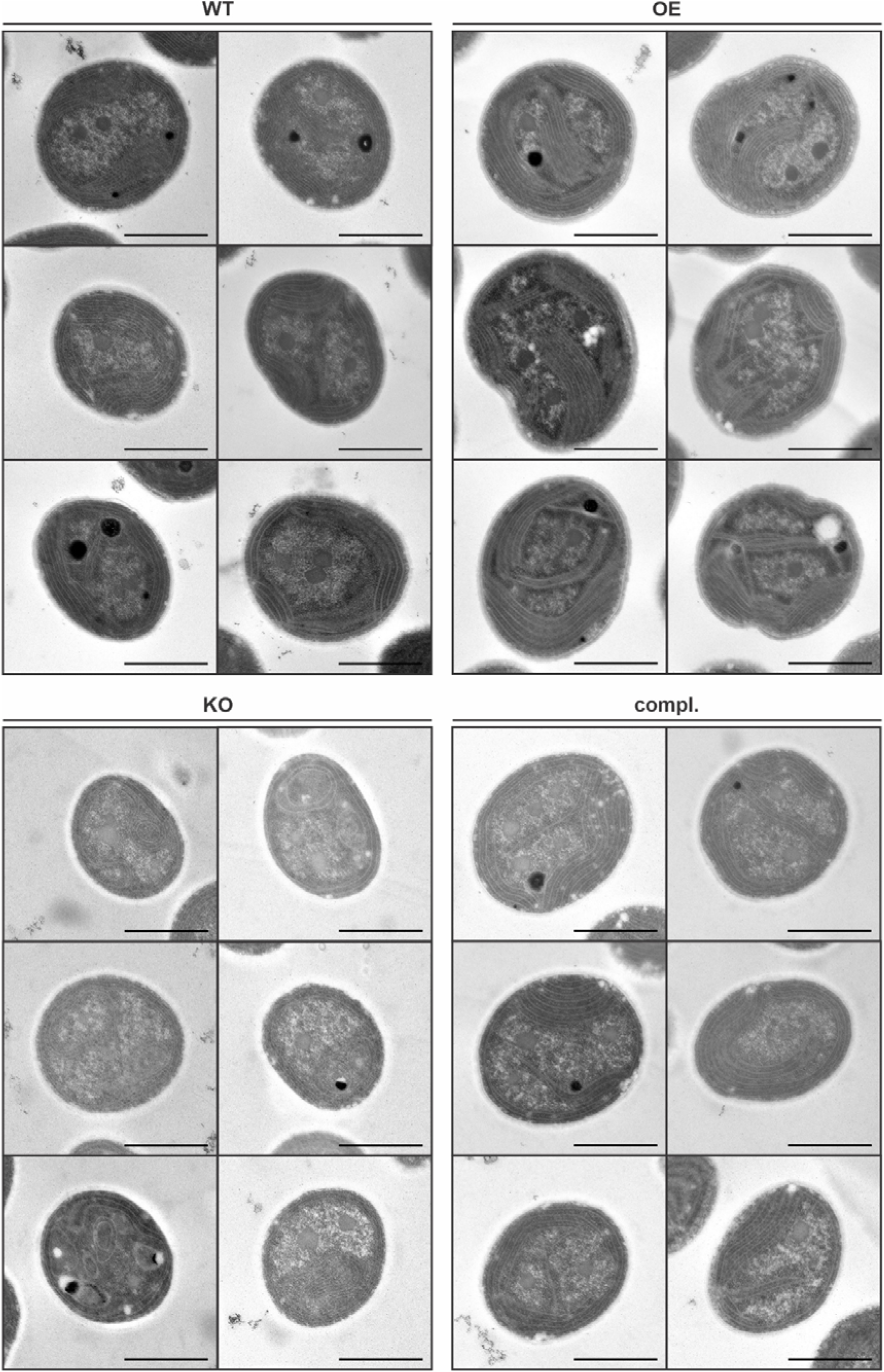
Transmission electron micrographs of *Synechocystis curT* expression level mutant cells. Representative cells of of *n* = 37/38/37/35 individual cells imaged for WT/OE/KO/compl., respectively, are shown. Scale bars indicate 1 µm.

**Supplementary Figure 4.**
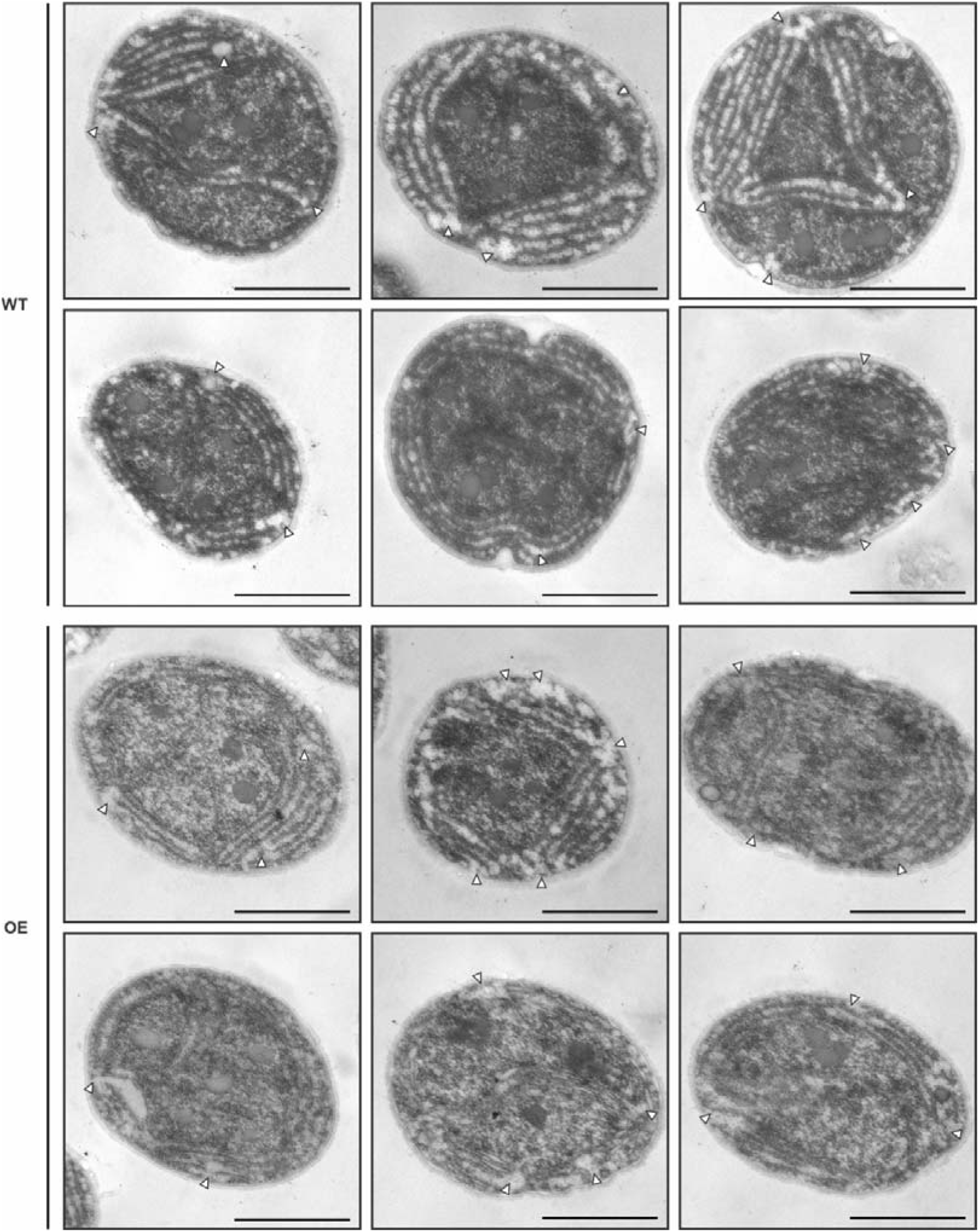
Transmission electron micrographs of *Synechocystis* WT and *curT* OE mutant cells grown under high light. Representative cells of of *n* = 20 individual cells imaged for WT/OE, respectively, are shown. White arrowheads show glycogen granula overlapping with tentative TCZs. Scale bars indicate 1 µm.

**Supplementary Figure 5.**
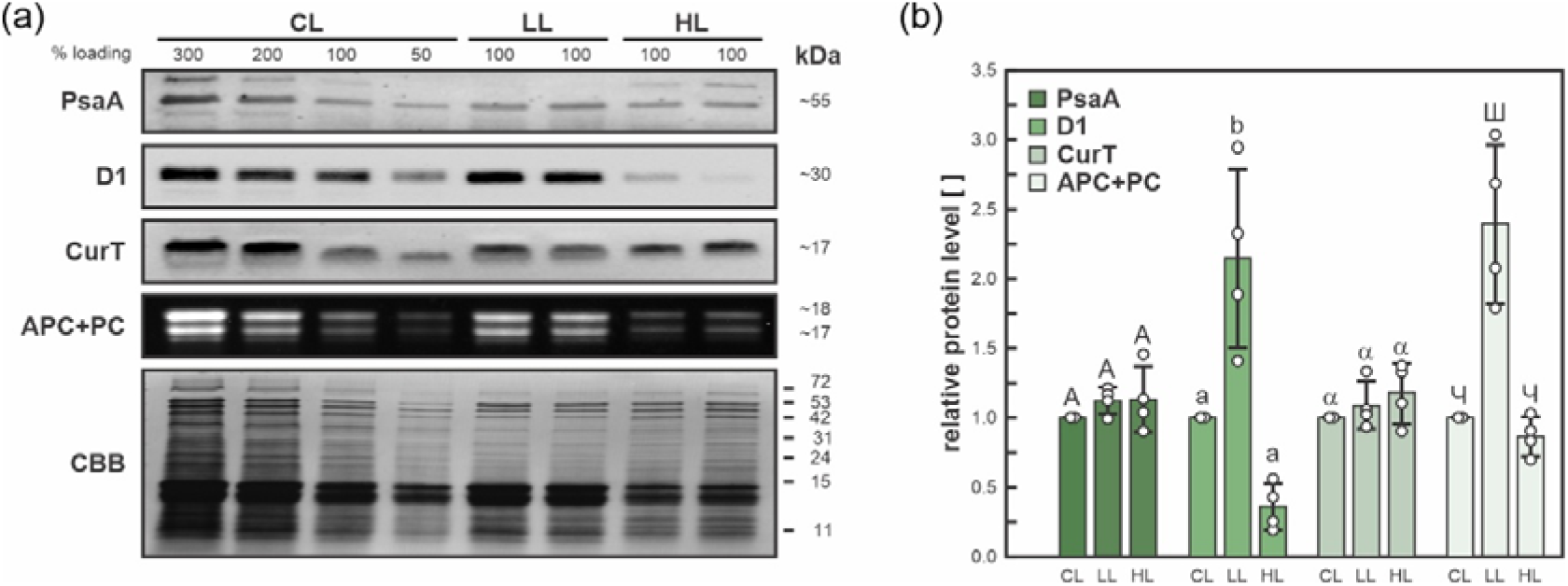
CurT protein level response to low and high light conditions. (a) PsaA, D1, and CurT protein levels as assessed by immunoblot analysis of whole-cell protein extracts obtained from mid-exponential phase cultures grown under control-(CL), low-(LL), and high-light (HL) conditions (50/20/300 µmol photons m^-2^ s^-1^ PAR, respectively) and harvested at 3-4 days past inoculation. Combined allophycocyanin and phycocyanin (APC+PC) levels were assessed by in-gel fluorescent assay prior to blotting of SDS-PA gels. Loading was adjusted to OD750. PVDF membrane Coomassie Brilliant Blue (CBB) staining served as loading control. Molecular weight markers are indicated. (b) Relative protein quantity estimates (CL 100% serving as reference; see Methods). Data shown represents *n* = 4 biological replicates. Columns show averages; error bars indicate standard deviations; individual data points are shown. Letters indicate statistically significant differences (*p*<0.05) according to *post-hoc* Bonferroni-Holm-corrected Tukey HSD after significant among-group differences (*p*<0.05) had been determined by two-sided one-way ANOVA.

**Supplementary Figure 6.**
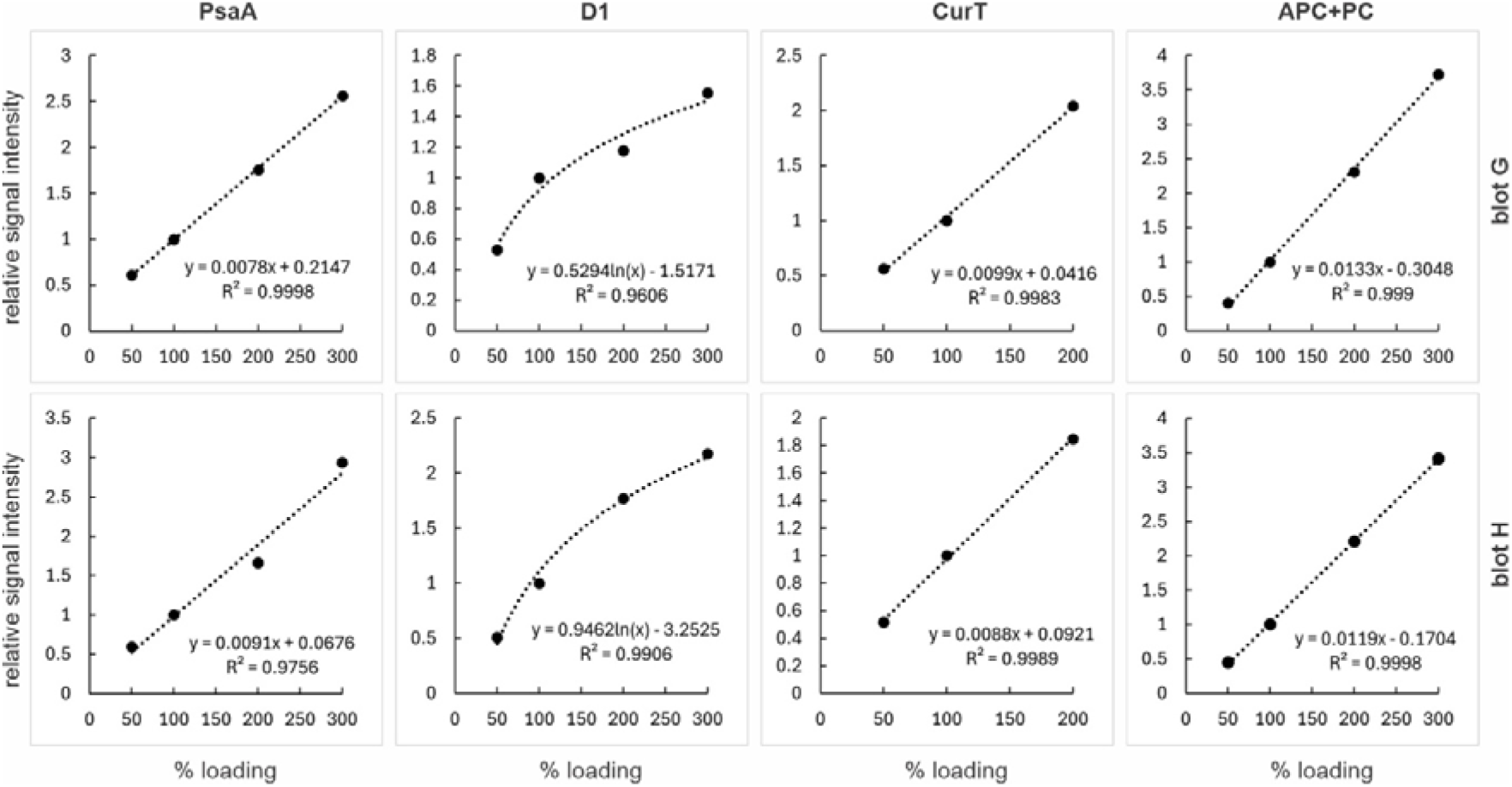
Calibration curves for *Synechocystis* WT control-/low-/high-light culture immunoblot signal quantification. Calibration curves derived from immunoblot ECL signals (PsaA, D1, CurT) and in-gel fluorescence signals prior to blotting (APC+PC) shown in Supplementary Fig. 5 (plus two more biological replicates; blots G-H) indicate partially linear (PsaA, CurT, APC+PC), partially logarithmic (D1) correlation of whole-cell protein extract sample input (x-axis; 100% corresponding to OD_750_=0.2 cell equivalents) and signal intensity relative to 100% control-light (CL) sample input (y-axis). Best-fitting linear or logarithmic regression models were chosen covering the required range for signal quantification of WT samples grown under low and high light (20 and 300 µmol photons m^-2^ s^-1^ PAR) relative to WT grown under CL conditions (50 µmol photons m^-2^ s^-1^ PAR). Calibration curve equations used for protein level approximation and corresponding coefficients of determination (R^2^) are indicated.

## References

Acuña,s A., Van Alphen, P., Van Grondelle, R., & Van Stokkum, I. (2018). The phycobilisome terminal emitter transfers its energy with a rate of (20 ps)–1 to photosystem II. Photosynthetica, 56(1), 265–274.

Armbruster, U., Labs, M., Pribil, M., Viola, S., Xu, W., Scharfenberg, M., Hertle, A. P., Rojahn, U., Jensen, P. E., Rappaport, F., Joliot, P., Dormann, P., Wanner, G., & Leister, D. (2013). Arabidopsis CURVATURE THYLAKOID1 proteins modify thylakoid architecture by inducing membrane curvature. Plant Cell, 25(7), 2661–2678. 10.1105/tpc.113.113118

Austin, J. R., & Staehelin, L. A. (2011). Three-dimensional architecture of grana and stroma thylakoids of higher plants as determined by electron tomography. Plant physiology, 155(4), 1601–1611.

Baikie, T. K., Wey, L. T., Lawrence, J. M., Medipally, H., Reisner, E., Nowaczyk, M. M., Friend, R. H., Howe, C. J., Schnedermann, C., & Rao, A. (2023). Photosynthesis re-wired on the pico-second timescale. Nature, 615(7954), 836–840.

Böde, K., Trotta, A., Dlouhý, O., Javornik, U., Paakkarinen, V., Fujii, H., Domonkos, I., Zsiros, O., Plavec, J., & Špunda, V. (2025). Lipid Phase Behaviour of the Curvature Region of Thylakoid Membranes of Spinacia oleracea. Physiologia plantarum, 177(3), e70289.

Choubeh, R. R., Wientjes, E., Struik, P. C., Kirilovsky, D., & Van Amerongen, H. (2018). State transitions in the cyanobacterium Synechococcus elongatus 7942 involve reversible quenching of the photosystem II core. Biochimica et Biophysica Acta (BBA)-Bioenergetics, 1859(10), 1059–1066.

Dann, M., Kim, E., Fujimura-Kamada, K., Berisha, V., Nomura, M., Pohland, A.-C., Watanabe, M., Ostermeier, M., Sommer, F., & Schroda, M. (2025). CurT/CURT1 proteins are involved in cell and chloroplast division coordination of cyanobacteria and green algae. Nature communications, 16(1), 8424.

Dann, M., Ortiz, E. M., Thomas, M., Guljamow, A., Lehmann, M., Schaefer, H., & Leister, D. (2021). Enhancing photosynthesis at high light levels by adaptive laboratory evolution. Nature plants, 7(5), 681–695.

de Jesus, A. J., Kastelowitz, N., & Yin, H. (2013). Changes in lipid density induce membrane curvature. RSC advances, 3(33), 13622–13625.

Dobakova, M., Tichy, M., & Komenda, J. (2007). Role of the PsbI protein in photosystem II assembly and repair in the cyanobacterium Synechocystis sp. PCC 6803. Plant Physiol, 145(4), 1681–1691. 10.1104/pp.107.107805

Espinoza-Corral, R., Iwai, M., Zavřel, T., Lechno-Yossef, S., Sutter, M., Červený, J., Niyogi, K. K., & Kerfeld, C. A. (2024). Phycobilisome protein ApcG interacts with PSII and regulates energy transfer in Synechocystis. Plant physiology, 194(3), 1383–1396.

Gandini, C., Schmidt, S. B., Husted, S., Schneider, A., & Leister, D. (2017). The transporter SynPAM71 is located in the plasma membrane and thylakoids, and mediates manganese tolerance in Synechocystis PCC6803. New Phytol, 215(1), 256–268. 10.1111/nph.14526

Garcia-Pichel, F., Zehr, J. P., Bhattacharya, D., & Pakrasi, H. B. (2020). What’s in a name? The case of cyanobacteria. Journal of phycology, 56(1), 1–5.

Georg, J., Voß, B., Scholz, I., Mitschke, J., Wilde, A., & Hess, W. R. (2009). Evidence for a major role of antisense RNAs in cyanobacterial gene regulation. Molecular Systems Biology, 5(1), 305.

Gu, L., Grodzinski, B., Han, J., Marie, T., Zhang, Y. J., Song, Y. C., & Sun, Y. (2022). Granal thylakoid structure and function: explaining an enduring mystery of higher plants. New Phytologist, 236(2), 319–329.

Heinz, S., Rast, A., Shao, L., Gutu, A., Gugel, I. L., Heyno, E., Labs, M., Rengstl, B., Viola, S., Nowaczyk, M. M., Leister, D., & Nickelsen, J. (2016). Thylakoid Membrane Architecture in Synechocystis Depends on CurT, a Homolog of the Granal CURVATURE THYLAKOID1 Proteins. Plant Cell, 28(9), 2238–2260. 10.1105/tpc.16.00491

Hong, S.-J., Kim, H., Jang, J. H., Cho, B.-K., Choi, H.-K., Lee, H., & Lee, C.-G. (2014). Proteomic analysis of Synechocystis sp. PCC6803 responses to low-temperature and high light conditions. Biotechnology and bioprocess engineering, 19(4), 629–640.

Huokko, T., Ni, T., Dykes, G. F., Simpson, D. M., Brownridge, P., Conradi, F. D., Beynon, R. J., Nixon, P. J., Mullineaux, C. W., & Zhang, P. (2021). Probing the biogenesis pathway and dynamics of thylakoid membranes. Nature communications, 12(1), 1–14.

Izuhara, T., Kaihatsu, I., Jimbo, H., Takaichi, S., & Nishiyama, Y. (2020). Elevated Levels of Specific Carotenoids During Acclimation to Strong Light Protect the Repair of Photosystem II in Synechocystis sp. PCC 6803. Front Plant Sci, 11, 1030. 10.3389/fpls.2020.01030

Jackson, P. J., Hitchcock, A., Brindley, A. A., Dickman, M. J., & Hunter, C. N. (2023). Absolute quantification of cellular levels of photosynthesis-related proteins in Synechocystis sp. PCC 6803. Photosynthesis Research, 155(3), 219–245.

Kılıç, M., Gollan, P. J., Lepistö, A., Isojärvi, J., Sakurai, I., Aro, E. M., & Mulo, P. (2022). Gene expression and organization of thylakoid protein complexes in the PSII-less mutant of Synechocystis sp. PCC 6803. Plant Direct, 6(6), e409.

Kirst, H., Gabilly, S. T., Niyogi, K. K., Lemaux, P. G., & Melis, A. (2017). Photosynthetic antenna engineering to improve crop yields. Planta, 245(5), 1009–1020.

Klinkert, B., Ossenbuhl, F., Sikorski, M., Berry, S., Eichacker, L., & Nickelsen, J. (2004). PratA, a periplasmic tetratricopeptide repeat protein involved in biogenesis of photosystem II in Synechocystis sp. PCC 6803. J Biol Chem, 279(43), 44639–44644. 10.1074/jbc.M405393200

Komenda, J., Sobotka, R., & Nixon, P. J. (2024). The biogenesis and maintenance of PSII: Recent advances and current challenges. The Plant Cell, koae082.

Koyama, K., Suzuki, H., Noguchi, T., Akimoto, S., Tsuchiya, T., & Mimuro, M. (2008). Oxygen evolution in the thylakoid-lacking cyanobacterium Gloeobacter violaceus PCC 7421. Biochimica et Biophysica Acta (BBA)-Bioenergetics, 1777(4), 369–378.

Kusama, Y., Inoue, S., Jimbo, H., Takaichi, S., Sonoike, K., Hihara, Y., & Nishiyama, Y. (2015). Zeaxanthin and Echinenone Protect the Repair of Photosystem II from Inhibition by Singlet Oxygen in Synechocystis sp. PCC 6803. Plant Cell Physiol, 56(5), 906–916. 10.1093/pcp/pcv018

Lea-Smith, D. J., Ortiz-Suarez, M. L., Lenn, T., Nürnberg, D. J., Baers, L. L., Davey, M. P., Parolini, L., Huber, R. G., Cotton, C. A., & Mastroianni, G. (2016). Hydrocarbons are essential for optimal cell size, division, and growth of cyanobacteria. Plant physiology, 172(3), 1928–1940.

Liberton, M., Howard Berg, R., Heuser, J., Roth, R., & Pakrasi, H. B. (2006). Ultrastructure of the membrane systems in the unicellular cyanobacterium Synechocystis sp. strain PCC 6803. Protoplasma, 227(2-4), 129–138. 10.1007/s00709-006-0145-7

Lübben, M. K., Klingl, A., Nickelsen, J., & Ostermeier, M. (2024). CLEM, a universal tool for analyzing structural organization in thylakoid membranes. Physiologia plantarum, 176(4), e14417.

Ma, L., Dong, B., Sun, M., Hao, R., Wang, X., Yu, H., Han, C., Muhire, A., Gachie, S. W., & Li, D. (2025). VESICLE-INDUCING PROTEIN IN PLASTIDS 1 from thylakoid-lacking Gloeobacter promotes thylakoid formation in Arabidopsis. Plant physiology, kiaf359.

Mahbub, M., Hemm, L., Yang, Y., Kaur, R., Carmen, H., Engl, C., Huokko, T., Riediger, M., Watanabe, S., Liu, L.-N., Wilde, A., Hess, W. R., & Mullineaux, C. W. (2020). mRNA localization, reaction centre biogenesis and thylakoid membrane targeting in cyanobacteria. Nature plants, 6(9), 1179–1191.

Mareš, J., Strunecký, O., Bučinská, L., & Wiedermannová, J. (2019). Evolutionary patterns of thylakoid architecture in cyanobacteria. Frontiers in Microbiology, 10, 277.

McCauley, S., & Melis, A. (1987). Quantitation of photosystem II activity in spinach chloroplasts. Effect of artificial quinone acceptors. Photochemistry and photobiology, 46(4), 543–550.

Murphy, D. J. (1982). The importance of non-planar bilayer regions in photosynthetic membranes and their stabilisation by galactolipids. FEBS letters, 150(1), 19–26.

Myers, J. A., Curtis, B. S., & Curtis, W. R. (2013). Improving accuracy of cell and chromophore concentration measurements using optical density. BMC biophysics, 6(1), 4.

Nickelsen, J., & Rengstl, B. (2013). Photosystem II assembly: from cyanobacteria to plants. Annual review of plant biology, 64, 609–635.

Ogawa, T., Misumi, M., & Sonoike, K. (2017). Estimation of photosynthesis in cyanobacteria by pulse-amplitude modulation chlorophyll fluorescence: problems and solutions. Photosynthesis Research, 133(1), 63–73.

Ostermeier, M., Buschmann, I., Heinz, S., & Nickelsen, J. (2025). The subcellular localisation of early photosystem II assembly in Synechocystis sp. PCC 6803. Physiologia plantarum, 177(2), e70234.

Ostermeier, M., Garibay-Hernández, A., Holzer, V. J., Schroda, M., & Nickelsen, J. (2024). Structure, biogenesis, and evolution of thylakoid membranes. The Plant Cell, 36(10), 4014–4035.

Ostermeier, M., Heinz, S., Hamm, J., Zabret, J., Rast, A., Klingl, A., Nowaczyk, M. M., & Nickelsen, J. (2022). Thylakoid attachment to the plasma membrane in Synechocystis sp. PCC 6803 requires the AncM protein. The Plant Cell, 34(1), 655–678.

Peschke, M., Moog, D., Klingl, A., Maier, U. G., & Hempel, F. (2013). Evidence for glycoprotein transport into complex plastids. Proceedings of the National Academy of Sciences, 110(26), 10860–10865.

Pribil, M., Sandoval-Ibáñez, O., Xu, W., Sharma, A., Labs, M., Liu, Q., Galgenmueller, C., Schneider, T., Wessels, M., & Matsubara, S. (2018). Fine-tuning of photosynthesis requires CURVATURE THYLAKOID1-mediated thylakoid plasticity. Plant physiology, 176(3), 2351–2364.

Qi, M., Zhao, Z., & Nixon, P. (2023). The photosynthetic electron transport chain of oxygenic photosynthesis. Bioelectricity 5 (1): 31–38. In.

Rast, A., Schaffer, M., Albert, S., Wan, W., Pfeffer, S., Beck, F., Plitzko, J. M., Nickelsen, J., & Engel, B. D. (2019). Biogenic regions of cyanobacterial thylakoids form contact sites with the plasma membrane. Nat Plants, 5(4), 436–446. 10.1038/s41477-019-0399-7

Rengstl, B., Oster, U., Stengel, A., & Nickelsen, J. (2011). An intermediate membrane subfraction in cyanobacteria is involved in an assembly network for Photosystem II biogenesis. J Biol Chem, 286(24), 21944–21951. 10.1074/jbc.M111.237867

Rexroth, S., Mullineaux, C. W., Ellinger, D., Sendtko, E., Rögner, M., & Koenig, F. (2011). The plasma membrane of the cyanobacterium Gloeobacter violaceus contains segregated bioenergetic domains. The Plant Cell, 23(6), 2379–2390.

Reynolds, E. S. (1963). The use of lead citrate at high pH as an electron-opaque stain in electron microscopy. The Journal of cell biology, 17(1), 208.

Rippka, R., Deruelles, J., Waterbury, J. B., Herdman, M., & Stanier, R. Y. (1979). Generic assignments, strain histories and properties of pure cultures of cyanobacteria. Microbiology, 111(1), 1–61.

Sacharz, J., Bryan, S. J., Yu, J., Burroughs, N. J., Spence, E. M., Nixon, P. J., & Mullineaux, C. W. (2015). Sub-cellular location of FtsH proteases in the cyanobacterium Synechocystis sp. PCC 6803 suggests localised PSII repair zones in the thylakoid membranes. Mol Microbiol, 96(3), 448–462. 10.1111/mmi.12940

Schägger, H., & Von Jagow, G. (1987). Tricine-sodium dodecyl sulfate-polyacrylamide gel electrophoresis for the separation of proteins in the range from 1 to 100 kDa. Analytical biochemistry, 166(2), 368–379.

Schneider, C. A., Rasband, W. S., & Eliceiri, K. W. (2012). NIH Image to ImageJ: 25 years of image analysis. Nature methods, 9(7), 671–675.

Selao, T. T., Zhang, L., Knoppova, J., Komenda, J., & Norling, B. (2016). Photosystem II Assembly Steps Take Place in the Thylakoid Membrane of the Cyanobacterium Synechocystis sp. PCC6803. Plant Cell Physiol, 57(1), 95–104. 10.1093/pcp/pcv178

Sengupta, A., Bandyopadhyay, A., Schubert, M. G., Church, G. M., & Pakrasi, H. B. (2023). Antenna modification in a fast-growing cyanobacterium Synechococcus elongatus UTEX 2973 leads to improved efficiency and carbon-neutral productivity. Microbiology spectrum, 11(4), e00500–00523.

Stengel, A., Gugel, I. L., Hilger, D., Rengstl, B., Jung, H., & Nickelsen, J. (2012). Initial steps of photosystem II de novo assembly and preloading with manganese take place in biogenesis centers in Synechocystis. Plant Cell, 24(2), 660–675. 10.1105/tpc.111.093914

Toyoshima, M., Sakata, M., Ueno, Y., Toya, Y., Matsuda, F., Akimoto, S., & Shimizu, H. (2021). Proteome analysis of response to different spectral light irradiation in Synechocystis sp. PCC 6803. Journal of Proteomics, 246, 104306.

van de Meene, A. M., Hohmann-Marriott, M. F., Vermaas, W. F., & Roberson, R. W. (2006). The three-dimensional structure of the cyanobacterium Synechocystis sp. PCC 6803. Arch Microbiol, 184(5), 259–270. 10.1007/s00203-005-0027-y

van Stokkum, I. H., Akhtar, P., Biswas, A., & Lambrev, P. H. (2023). Energy transfer from phycobilisomes to photosystem I at 77 K. Frontiers in plant science, 14, 1293813.

Vanni, S., Hirose, H., Barelli, H., Antonny, B., & Gautier, R. (2014). A sub-nanometre view of how membrane curvature and composition modulate lipid packing and protein recruitment. Nature communications, 5(1), 4916.

Walther, P., & Ziegler, A. (2002). Freeze substitution of high-pressure frozen samples: the visibility of biological membranes is improved when the substitution medium contains water. Journal of microscopy, 208(1), 3–10.

Zavřel, T., Sinetova, M. A., & Červený, J. (2015). Measurement of chlorophyll a and carotenoids concentration in cyanobacteria. Bio-protocol, 5(9), e1467–e1467.

Zhang, L., Selao, T. T., Selstam, E., & Norling, B. (2015). Subcellular Localization of Carotenoid Biosynthesis in Synechocystis sp. PCC 6803. PLoS One, 10(6), e0130904. 10.1371/journal.pone.0130904

Zhang, Z., Ge, X., Huokko, T., & Liu, L. N. (2025). Curvature Thylakoid1-like protein CurT mediates thylakoid membrane architecture in Synechococcus elongatus PCC 7942. Mlife, 4(5), 567–571.

